# SureQuant™ IS-PRM Enables Cross-Species Targeted Quantification of Retinoid Metabolism and Signaling Proteins in the Heart

**DOI:** 10.64898/2026.07.27.741027

**Authors:** Sogol Sedighi, Robert N. O’Meally, Sunayana Begum Syed, Hugo Amadei, Biyi Li, Chulan Kwon, Abhishek Vats, Don Zack, Kenneth B. Margulies, Brian O’ Rourke, Peter M. Abadir, Robert N. Cole, D. Brian Foster

## Abstract

Retinoic acid signaling is critical for cardiac development and homeostasis. Dysregulation of all-trans retinoic acid metabolism contributes to vascular atherogenesis, restenosis, calcification, and heart failure. Therefore, assessment of proteins involved in retinoid metabolism and signaling has gained interest for identifying potential biomarkers and therapeutic targets in cardiovascular disease. However, quantifying these proteins remains challenging due to limitations of antibody-based methods. We developed a targeted proteomics approach using SureQuant™ internal standard-triggered parallel reaction monitoring mass spectrometry to profile these proteins. We designed a panel of 80 stable isotope-labeled (heavy) peptides representing proteins involved in retinoid signaling and metabolism, with sequences applicable to human samples and conserved across multiple species. Survey experiments using directed data-dependent acquisition on the Orbitrap Fusion Lumos mass spectrometer determined precursor and product ion masses for each heavy peptide, which were programmed into the SureQuant method for continuous monitoring. Upon detection of these heavy internal standards, the instrument transitions to a targeted PRM acquisition mode in which repeated high-resolution MS/MS spectra of both endogenous (light) and heavy peptides are acquired. Using this method, retinoid pathway-associated proteins were quantified to as low as 10 attomoles for selected targets across multiple tissues and developmental stages. Distinct tissue-specific retinoid metabolic networks were identified across lung, liver, retinal cell lines and cardiac tissues. Developmental profiling of mouse and rat hearts revealed remodeling of retinoid pathway proteins from embryonic to postnatal and adult stages, suggesting a functional transition from retinoid-driven cardiac development toward maintenance of retinoid homeostasis in the mature heart.

## Introduction

All-trans retinoic acid (ATRA) is the bioactive metabolite of vitamin A (retinol). It functions as a potent and pleiotropic regulator of gene expression through activation of nuclear retinoic acid receptors, influencing a wide range of developmental processes(1). Precise regulation of retinoic acid (RA) signaling is particularly critical during cardiac development, where both deficiency and excess are associated with severe congenital abnormalities (2). Beyond development, dysregulated retinoid homeostasis has also been implicated in adult cardiovascular disease. Low circulating RA levels are associated with an increased risk of all-cause cardiovascular mortality(3). In idiopathic dilated cardiomyopathy, we and others have shown that intracardiac ATRA levels are markedly reduced despite preserved retinol stores, suggesting impaired local retinoid metabolism rather than systemic deficiency. Similar reductions in cardiac ATRA precede functional decompensation in experimental pressure-overload heart failure, and restoration of myocardial ATRA levels can prevent disease progression(4). Despite its biological importance, direct therapeutic manipulation of ATRA signaling is challenging due to the pleiotropic effects of retinoids and the short half-life of ATRA in circulation. An alternative strategy is to target the enzymes that manage retinoid metabolism within specific tissues.

Retinol is obtained directly from the consumption of eggs, dairy, liver, and fish, or synthesized in the intestine from β-carotene (5–9) found in vibrantly colored vegetables (10–13). Following intestinal absorption, retinol is transported via the circulation to the liver, where it is stored as retinyl esters or used to fine tune hepatic glucose and fatty acid metabolism (14, 15). Retinol stores can be mobilized from the liver to ensure adequate distribution as free retinol to peripheral tissues where it is stored locally as a ready reserve for in situ ATRA production. Each step in the conversion of retinol to retinaldehyde to retinoic acid, and its downstream inactive polar metabolites, is catalyzed by different families of enzymes with multiple isoforms, namely specific families of retinol dehydrogenases (Rdh), retinal reductases (Rrd, Dhrs), retinaldehyde dehydrogenases (Raldh, Aldh1a), and p450 hydroxylases (Cyp26)(3). The combination of these enzymes varies by cell-type within a tissue to yield context-specific function and regulation. For instance, the enzymes of retinoid metabolism vary between hepatic stellate cells and hepatocytes, and differ again from those of epidermal cells and adipocytes. Thus each organ, or more specifically, each cell type within an organ, metabolize retinol differently(16, 17).

Decoding a cell’s retinoid metabolic and signaling program systematically is challenging, owing to the technical limitations of antibody-based assays. While Western blotting is widely used, it is inherently low throughput, limited in multiplexing capacity, and dependent on the availability and quality of commercially available antibodies(18, 19). Targeted mass spectrometry technology for quantifying relevant predefined targets, such as parallel reaction monitoring (PRM), provides for sensitive and protein-specific peptide detection without antibodies. Moreover, PRM can be scaled to detect multiple proteins of interest including metabolic and signaling pathways. However, large-scale multiplexing is challenging because it relies on scheduled acquisition windows that are highly sensitive to minor chromatographic drifts.

Internal standard-triggered PRM (IS-PRM) uses the detection of spiked-in stable-isotope-labeled internal standards, rather than retention time, as the cue to trigger PRM, enabling the reliable detection of even larger numbers of targets without compromising data quality or measurement sensitivity(20, 21). The Thermo Scientific™ SureQuant™ IS Targeted Quantitation workflow builds on the original IS-PRM approach, that streamlines creating an IS-PRM assay, simplifying the process and enhancing its usability(20).

In this study, we developed a SureQuant-based targeted proteomic assay for the retinoid signaling pathway that is broadly applicable across diverse cell types and species. Peptides were selected primarily for targeted quantification of human proteins, while also maximizing compatibility with commonly used preclinical animal models. We aimed to establish a robust method for characterizing retinoid signaling proteins across different cell lysates, tissues, and stages of heart development.

## Materials and Methods

### Human Myocardial Tissue

Human myocardial tissue was obtained under protocols approved by the Institutional Review Boards of the University of Pennsylvania and the Gift-of-Life Donor Program (Pennsylvania, USA). Hearts were collected from cadaveric donors at the time of organ donation. In all cases, hearts were arrested in situ using ice-cold cardioplegia solution and transported on wet ice. Whole hearts and dissected left ventricular cavities were weighed to assess the degree of hypertrophy. Transmural myocardial samples were obtained from the mid–left ventricular free wall below the papillary muscle(22).

### Cell Line Lysates and Organoid Samples

Commercially available whole-cell lysates were used for selected experiments, including A549 cells (Abcam, AB7910-1001), HepG2 cells (Abcam, ab166833-1002), and human retinal pigment epithelial cell lysates (ScienCell, #6546). Retinal organoids (RO) were generated using modified H9 embryonic stem cells (H9-ESC) (23) as described before (24). Briefly, H9-ESC was de-adhered using accumax and 90 K cells/well were seeded in the AggreWell^™^800 24-well plate (Stem cells, #34815) in 1.5 mL mTeSR™ Plus with 10 µM blebistatin (Sigma, #B0560). mTeSR plus medium was replaced by 500 µL, 750 µL, and 1.5 mL with neural induction media (NIM), every 24 hours followed by maintenance in the NIM. The embryoid bodies were transferred to Matrigel coated plate for optic vesicle formation in NIM medium followed by replacing 3:1 medium after day 16. RO were scraped at day 28 to maintain in the suspension in a low attachment plate. The groomed RO were switched to 3:1+Taurine+FBS medium till day 70 followed by addition of 1 µM RA till day 98 and then switched to N2 medium with 0.5 µM RA for photoreceptors maturation till 300 days. ROs at day 294 were collected for this experiment.

### Animal Models

All animal studies were approved by the Institutional Animal Care and Use Committee (IACUC) at Johns Hopkins University and conducted in accordance with AVMA guidelines. Animals were housed under specific pathogen-free conditions with controlled temperature and humidity, a 12-hour light/dark cycle, and access to food and water. Mice were euthanized by isoflurane overdose, after which hearts were rapidly excised following thoracotomy, rinsed in ice-cold PBS, and processed for downstream analyses, including snap-freezing in liquid nitrogen. Fresh instruments were used for each animal to prevent cross-contamination. Rat pup hearts were collected at postnatal days 0 and 14 from pups, with three biological replicates per time point. Mouse embryonic hearts were harvested at embryonic day 9.5 (E9.5) from timed-pregnant C57BL/6J mice (Jackson Laboratory, strain 000664) by micro-dissecting the linear heart tube together with the adjacent dorsal trunk tissue containing cardiac progenitor cells. Neonatal mouse hearts were collected at postnatal day 1 (P1), and adult hearts were collected from 12-week-old male C57BL/6J mice (Jackson Laboratory, strain 000664).

### Immunoblotting

Protein extracts were solubilized and denatured in 1× LDS sample buffer and separated by SDS-PAGE using 4–12% NuPAGE Bis-Tris gels (1 mm; Invitrogen). Electrophoresis was performed at room temperature for 35 min at 200 V. Proteins were transferred to nitrocellulose membranes using a Trans-Blot Turbo Transfer System (Bio-Rad). Following transfer, membranes were stained with Revert 700 Total Protein Stain (LI-COR Biosciences) to assess transfer efficiency and total protein loading. Membranes were blocked for 1 h at room temperature in Tris-buffered saline (TBS) containing 2.5% bovine serum albumin (BSA) and 0.1% Tween-20 and subsequently incubated overnight at 4°C with primary antibodies diluted in blocking buffer. Following washing with TBS supplemented with 0.1% (v/v) Tween-20 (TBST), membranes were incubated for 1 h at room temperature with IRDye-conjugated secondary antibodies diluted in blocking buffer (IRDye 800CW Donkey Anti-Rabbit IgG, IRDye 680LT Donkey Anti-Mouse IgG (LI-COR Biosciences)). Fluorescence signals were detected using an Odyssey infrared imaging system, and band intensities were quantified using Odyssey Application Software version 3.0 (LI-COR Biosciences). Densitometric analysis was performed using ImageJ software on digitized TIFF image files. Protein abundance was normalized to total protein levels determined by Revert 700 Total Protein Stain. Primary antibodies used included DHRS4 (Proteintech, Cat. No. 15279-1-AP), and ALDH1A2 (Abcam, Cat. No. ab156019), RBP1 (Proteintech, Cat. No 22683-1-AP).

### Selection of Peptides as Standards

A publicly available spectral library of retinoid-associated proteins from MassIVE (Mass Spectrometry Interactive Virtual Environment) was imported into Skyline (MacCoss Lab Software (25)), and peptides were ranked based on their peak intensities. Peptides containing cysteine (Cys), methionine (Met), or histidine (His), as well as peptides harboring predicted N-linked glycosylation motifs (NXT/NXS), arginine–proline (RP) or lysine–proline (KP) motifs were excluded from further consideration. A final panel of **80** tryptic peptides representing **49** retinoid-associated proteins was curated from the initial candidate list based on multiple criteria, including reproducibility, sensitivity, optimal spectral library matching in Skyline, cross-species applicability, with a primary focus on peptide sequence identity between *Homo sapiens*, *Rattus norvegicus*, and *Mus musculus*. Tryptic peptides were screened across species using the MaCPepDB database (25). For the selected peptide panel, stable isotope–labeled (“heavy”) peptides incorporating ^13^C_6_^15^N_2_ labels at the C-terminal lysine or arginine residues (K+8 and R+10) were synthesized and ordered from JPT Peptide Technologies (Berlin, Germany). Corresponding unlabeled (“light”) synthetic peptides were also obtained from JPT Peptide Technologies.

### Sample Preparation and Digestion

Heart tissues were homogenized using a handheld Polytron tissue disrupter in filtered, deionized Tris-buffered 9 M urea (5 mL, pH 7.5) supplemented with protease and phosphatase inhibitors (1×; cOmplete™, Mini, EDTA-free, Roche; PhosSTOP™, Roche) and allowed to solubilize for 30 min at room temperature. Retinal organoids (ROs) were washed with PBS and lysed in RIPA buffer containing the same inhibitors, 5% SDS, and 1 mM DTT. All samples, including the cell lysates, lysed heart tissues, and lysed retinal organoids, were subjected to methanol/chloroform/water extraction and protein precipitation by the method of Wessel & Flugge (26). Samples were dried under nitrogen gas to remove residual chloroform before resolubilizing for 30 min in 9M urea. Aggregates were disrupted by brief bursts of sonication (< 30s total). 50 fmol of the “heavy” isotope-labeled peptides per each 5 ug of protein were added to each sample, and they all were subjected to proteolytic digestion with proteomics grade Trypsin/Lys-C (1 μg/100μg protein; Promega) at room temperature, overnight. The following morning, samples were supplemented with Trypsin/Lys-C (1 μg/100μg protein; Promega). At that time, DTT was added to samples at a concentration of 5 mM and the digest was allowed to proceed for 1 hour prior to peptide alkylation by addition of iodoacetamide to a final concentration of 15 mM. Alkylation was allowed to proceed for 30 minutes at room temperature in the dark. Peptides were subsequently acidified by the addition of trifluoroacetic acid to a final concentration 0.5% (v/v) and purified by solid phase extraction using SepPak tC18 cartridges (Waters) on a vacuum manifold. Purified peptides were eluted with 60% (v/v) acetonitrile in aqueous 0.1% (v/v) formic acid. Peptides were evaporated to dryness on an Eppendorf Vacufuge.

### LC/MS-MS analysis

Samples for SureQuant were analyzed using an Orbitrap Fusion Lumos mass spectrometer (Thermo Scientific) coupled with a Vanquish Neo nanoLC system (Thermo Scientific) Nanospray Flex ion source (Thermo Scientific), A 10-uL injection volume of sample was loaded by a trap and elute setup onto a 25 cm house packed 75 um ID fused silica nano column (Dr. Maisch 100A, 2.4um ReproSil phase). Peptides were eluted at a flow rate of 300 nL/minute across an optimized gradient consisting of 0.1% formic acid (buffer A) and 95% acetonitrile in 0.1% formic acid (buffer B) over 60 minutes. The eluted peptides were ionized at 2.2kV post column using a PepSep sprayer and 30um stainless steel emitter liquid junction setup (Bruker) that was coupled to the column using a Direct-Connect 360um FS to 1/16th fitting (VICI).

### Survey MS analyses

Prior to SureQuant acquisition, a survey of peptides by directed data-dependent acquisition (dDDA) was performed on an Orbitrap Fusion Lumos mass spectrometer (Thermo Scientific) using a mixture of heavy-labeled peptides spiked at 50 fmol per peptide. This was set up using a SureQuant Custom Panel Survey template in the Orbitrap Fusion Lumos software version 4.1. The resulting raw files were imported into Skyline using the standard Import Results workflow to evaluate heavy peptide detectability, confirm optimal precursor charge states and representative fragment ions, and to establish triggering thresholds (set to 1% of apex intensity of DDA survey analysis) for MS/MS scan initiation in subsequent SureQuant analyses.

### Targeted MS SureQuant analyses

The custom acquisition template available in Thermo Orbitrap Fusion Lumos software version 4.1 was used to develop the SureQuant assay. To implement this acquisition strategy, first the optimal precursor charge state, precursor intensity response, and diagnostic fragment ions for each peptide from survey MS analysis were incorporated into the SureQuant method, enabling targeted monitoring of the isotopically labeled trigger peptides. Standard MS parameters for SureQuant acquisition were as follows: spray voltage: 2.2kV, no sheath or auxiliary gas flow, heated capillary temperature 250 C. Full-scan mass spectra were acquired over a scan range of 300–1,500 m/z at a resolution of 120,000, with an AGC target of 300% corresponding to an absolute AGC value of 1.2 × 10⁶, and a maximum injection time (IT) of 50 ms. Within a 7-s cycle time following each MS1 scan, heavy peptides matching the expected m/z within 10 ppm and intensity threshold defined on the inclusion list were isolated (isolation width of 1.0 m/z) and fragmented (nCE:32) by HCD with a scan range: 100–1,700 m/z, maximum IT 20 ms, and normalized AGC target value: 1,000% (1e5), resolution: 15,000. A product ion trigger filter next performs pseudo-spectral matching, only triggering an MS/MS event of the endogenous, target peptide at the defined mass offset if n ≥ 4 product ions are detected from the defined list at their threshold value. If triggered, the subsequent light peptide MS/MS is scanned together with the heavy peptide in a PRM-type scan running at 60,000 resolutions. The maximum AGC for this scan type is set to 1000, and the maximum injection time is 116 msec for both heavy and light. The PRM type scan continues across the peaks until the threshold in the first trigger scan is no longer met, and the instrument returns to looking for trigger peptides.

### Calibration Curves with “heavy” and “light” peptides

Calibration curves were generated by spiking light synthetic peptides at 10, 25, 50, 100, 500, and 2000 attomoles on-column in the presence of a fixed 50 femtomoles of heavy internal standard peptides. For each peptide and concentration level, light and heavy peak areas were extracted. A correction factor was applied based on the ratio of the observed heavy signal to the median heavy signal across assays, and both heavy and light areas were scaled accordingly. Replicate measurements exhibiting high variability at individual concentration levels were excluded. Replicate measurements were summarized per peptide and concentration as mean corrected light area ± standard deviation (SD). The light/heavy ratio dot product (rdotp), exported from Skyline, was used as a measure of similarity between the endogenous peptide transition peak-area pattern and the corresponding heavy internal standard transition peak-area pattern(27, 28).

Linear regression was used to evaluate response linearity across the dynamic range, and per-peptide calibration plots were generated showing mean ± SD with the fitted regression line and corresponding R^2^ and p-values.

The limit of detection (LOD) was defined as the lowest peptide abundance at which the mean signal exceeded the mean blank signal plus three standard deviations of the blank (mean_blank + 3SD_blank), measured in a matrix of digested Saccharomyces cerevisiae protein. The limit of quantification (LOQ) was defined as the lowest peptide abundance at which the mean signal exceeded the mean blank signal plus ten standard deviations of the blank (mean_blank + 10SD_blank) in the same matrix. A practical limit of quantification (PLQ) was defined as the minimum abundance on the calibration curve for which the dot product between light and heavy peptides (rdotp) exceeded 0.7.

Analytical precision was evaluated by calculating the coefficient of variation (CV) from median light-to-heavy ratios across three replicate measurements in 2 fmol of injected light peptide. CV was calculated as the standard deviation divided by the mean of three replicate measurements.

### Data analyses

All MS data were imported into Skyline using the Import Results function. In Skyline, the MS/MS filtering acquisition method was set to SureQuant under the Full-Scan tab of the Transition Settings menu, and triggered chromatogram acquisition was enabled under the Instrument tab. The Skyline document generated during the survey setup, containing the curated heavy peptide panel, was used as a template. Endogenous (light) peptide counterparts corresponding to each heavy-labeled peptide were then generated within the Skyline document for targeted quantification. Skyline reports containing fragment ion peak areas for light and heavy peptides were exported and processed in R (v4.4.0). Quality filtering retained peptides with non-zero integrated peak area in at least three fragment ions for both heavy and endogenous peptides, and rdotp > 0.7 in at least one sample. Light-to-heavy ratios were calculated using the three most intense fragment ion transitions per peptide, computed as Σlight/Σheavy. A replicate-level normalization was applied in which the median heavy peptide signal from each injection was scaled to the experiment-wide median across replicates, and light-to-heavy ratios were adjusted accordingly. Endogenous peptide amounts were obtained by scaling normalized ratios to the known heavy peptide input (50 fmol per 5 µg total protein) and reported as attomoles per milligram of total protein. For heat map generation, data were log^2^-transformed, median-normalized, and row-wise z-scored. Heat maps were generated in R using the ComplexHeatmap package(29, 30). Hierarchical clustering of proteins was performed using complete linkage based on Euclidean distances. For clustering purposes only, missing values corresponding to signals below the limit of detection were temporarily imputed with low values to stabilize distance calculations; however, the heat maps display the original data, with missing values shown in light gray. Colors represent row-wise relative protein abundance across samples rather than raw abundances. For the PCA biplot, the PCAtools package was used on data that were log-transformed and median-normalized(31).

## Results

### Overview of SureQuant Workflow

As a trigger-based method, SureQuant depends on the precise detection of internal standard peptides to initiate targeted MS/MS acquisition of their endogenous counterparts(32). During SureQuant assay, the instrument first performs a high-resolution MS1 scan to monitor the heavy internal-standard (IS) peptides. When a heavy peptide is detected at the correct m/z and above a predefined intensity threshold, a fast, low-resolution MS/MS scan of the heavy peptide is acquired. If the expected fragment ions are confirmed, the instrument transitions into a targeted PRM acquisition mode in which repeated high-resolution MS/MS spectra of both the endogenous (light) and heavy peptide are acquired across the chromatogram until the threshold is no longer met, after which the instrument returns to “watch mode” to rapidly scan for other trigger peptides. (33) The trigger threshold therefore serves as both the start and stop signal for acquisition, such that the higher-resolution scans are only acquired while peptides of interest are actively eluting. This improves the duty cycle compared to scheduled PRM methods, which must continue acquiring across a broader retention time window regardless of whether the peptide of interest is eluting. **(Figure 1A).** Quantification is subsequently performed using the integrated fragment-ion peak areas derived from the acquired MS2 spectra. Following the dDDA survey assay **(Figure 1B-C),** all precursor intensity thresholds and fragment ion selections were obtained and used to develop the SureQuant method as described in the methods section. A custom acquisition template was structured such that the acquisition parameters for each unique isotopically labeled amino acid and charge state, are contained within a distinct branch originating from the full-scan node. In this study, four branches were used corresponding to R^2^⁺, R³⁺, K^2^⁺, and K³⁺ **(Figure 1D)**. During the initial survey scan, the precursor charge state yielding the highest precursor intensity and most consistent fragment ion quality was determined for each heavy synthetic peptide. Charge state selection was based on signal-to-noise ratio, fragment ion coelution, chromatographic peak shape, and spectral similarity (dotp) in Skyline. Peptides were then assigned to one of the four branches based on both the isotopically labeled amino acid they contained (heavy lysine(K) or heavy arginine(R)) and their optimal charge state. Within the Targeted Mass node, the optimal precursor intensity response, set to 1% of the maximum precursor height from the dDDA survey scan, was used to define the triggering threshold. Within the Targeted Mass Trigger node, fragment ions corresponding to the selected precursor were specified to enable spectral matching and subsequent triggering of PRM acquisition for peptide quantification.

**Figure 1.**
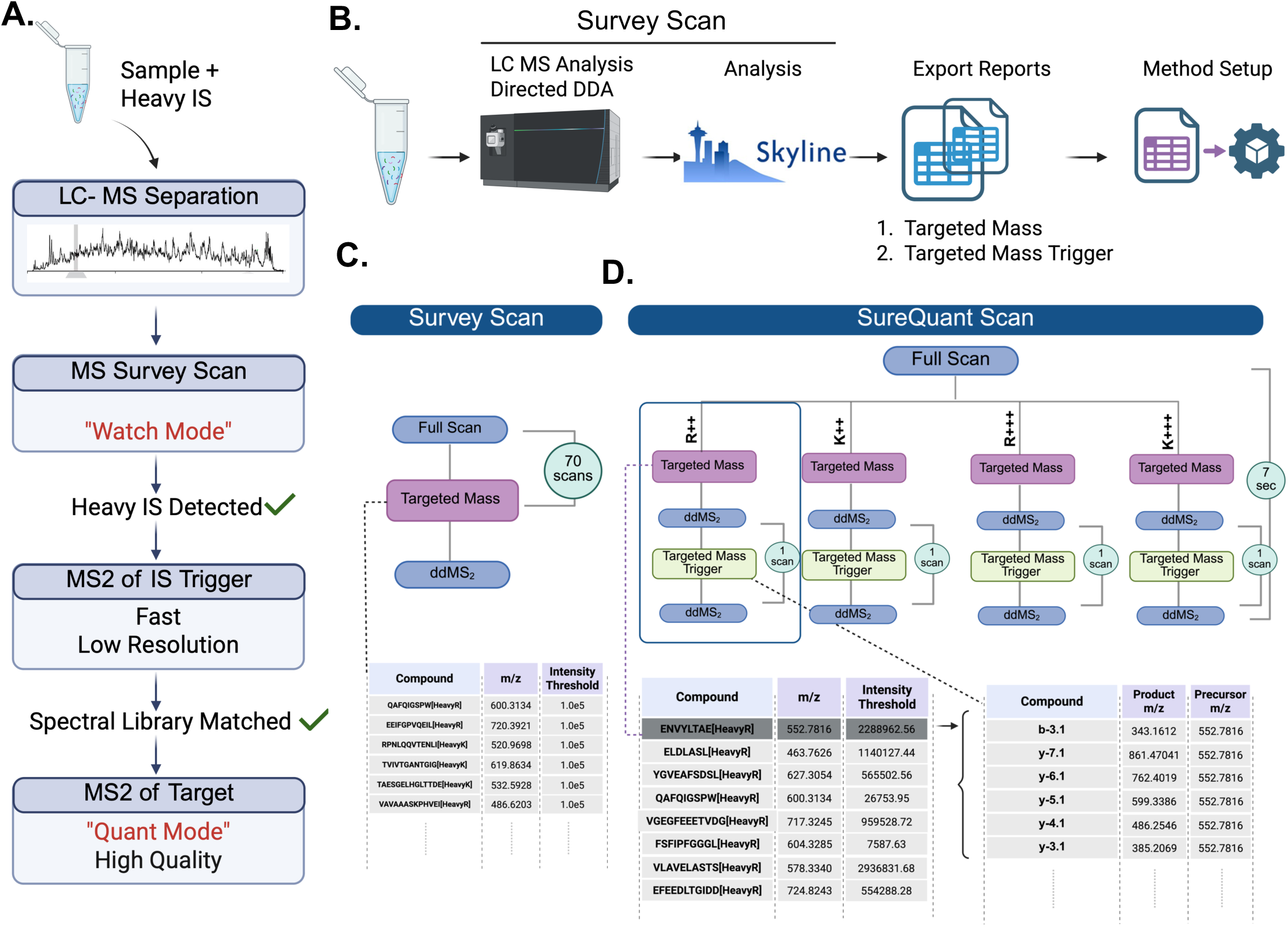
SureQuant acquisition workflow and Method Setup. **(A)** After LC separation of sample containing both light and heavy internal standard (IS), a full MS survey scan is performed in “watch mode.” When the heavy-labeled IS is detected, a rapid low-resolution MS^2^ scan is triggered for confirmation, followed by spectral-library matching. A subsequent high-resolution MS^2^ (“Quant Mode”) is then acquired for the corresponding endogenous (light) and heavy peptide.**(B)** For method setup preliminary survey scan is performed on heavy peptides using LC–MS/MS in data-dependent acquisition (DDA) mode. Representative fragment ions for each heavy peptide were selected, and precursor intensity thresholds were set at 1% of the apex intensity to enable MS/MS triggering in subsequent SureQuant analyses. These parameters were then exported to generate the targeted mass and targeted mass trigger inclusion lists used for instrument method configuration. **(C)** Within the Survey Scan node in Orbitrap Fusion Lumos, precursor inclusion entries containing the corresponding heavy peptide precursor m/z values were imported. A uniform precursor intensity threshold of 1.0 × 10⁵ was applied to all peptides **(D)** SureQuant node in Orbitrap Fusion Lumos software version 4.1 method editor. Each branch corresponds to a specific heavy isotope label and optimal precursor charge state. Within each branch, the Targeted Mass sub-node contains the precursor m/z and an intensity threshold set to 1% of the maximum precursor apex height calculated from the survey scan. Within the Targeted Mass Trigger sub-node, compound entries specify the fragment ion information, including fragment ion type, fragment ion ordinal, and product ion charge state (formatted as fragment ion type – fragment ion ordinal. product charge; e.g., b-3.1), along with product m/z and the corresponding precursor m/z.

### Cross-species applicability of the retinoid SureQuant panel

Standard peptides were selected for the design of targeted human protein assays, but with the additional goal of maximizing utility across commonly used preclinical animal models. Of the 80 peptides in the panel, 78 (98%) consisted of *Homo sapiens* (Hs) protein sequences, of which 40 (50%) had conserved sequence identity in *Rattus norvegicus* (Rn), while (41) (51%) peptides shared sequence identity in *Mus Musculus* (Mm). The panel also consisted of two Mm-Rn specific peptides **(Table 1)**.

**Table 1.** High Performing Tryptic Peptides Associated with Retinoid Mtabolism and Signaling.

| Table 1. High-Performing Tryptic Peptides Associated with Retinoid Metabolism and Signaling |  |  |  |  |  |  |  |
| --- | --- | --- | --- | --- | --- | --- | --- |
| Peptide | Protein | Name | LOD<sup>1</sup> | LOQ<sup>2</sup> | rdotpLOQ | PLQ<sup>3</sup> | Species<sup>5</sup> |
| EEIFGPVQEILR | ALDH1A2 | Aldehyde Dehydrogenase 1 Family Member A2 | 10 | 10 | 0.81 | 10 | Hs, Mm, Rn, Cp, Bt, Ss, Pt, Pa |
| FSFIPFGGGLR | CYP26A1 | Cytochrome P450 Family 26 Subfamily A Member 1 | 10 | 10 | 0.82 | 10 | Hs, Mm, Rn, Cp,Bt, Ss, Pt, Pa |
| LAQDGAHVVVSSR | DHRS4 | Dehydrogenase/Reductase SDR Family Member 4 | 10 | 10 | 0.71 | 10 | Hs, Bt, Ss, Pa |
| EEVEEVLDSK | RAI1 | Retinoic Acid Induced 1 | 10 | 10 | 0.82 | 10 | Hs, Pt |
| SLATWENENK | RBP1 | Retinol Binding Protein 1 | 10 | 10 | 0.82 | 10 | Hs, Cp, Ss,Pt, Pa |
| EFEEDLTGIDDR | RBP1 | Retinol Binding Protein 1 | 10 | 10 | 0.89 | 10 | Hs, Mm, Rn, Cp, Bt, Ss, Pt, Pa |
| VGEEFDEDNR | RBP7 | Retinol Binding Protein 7 | 10 | 10 | 0.93 | 10 | Hs, Pt, Pa |
| ENVYLTAER | RDH10 | Retinol Dehydrogenase 10 | 10 | 10 | 0.73 | 10 | Hs, Mm, Rn, Bt, Ss |
| IPFHDLQSEK | RDH12 | Retinol Dehydrogenase 12 | 10 | 10 | 0.82 | 10 | Hs, Pt, Pa |
| LFPQLEGK | RETSAT | Retinol Saturase | 10 | 10 | 0.74 | 10 | Hs, Mm, Rn, Cp, Ss, Pt, Pa |
| TVYSQSDLIASK | STRA8 | Stimulated by Retinoic Acid 8 | 10 | 10 | 0.78 | 10 | Hs, Pt |
| AVEAAQVAFQR | ALDH1A3 | Aldehyde Dehydrogenase 1 Family Member A3 | 10 | 25 | 0.73 | 25 | Hs, Pt |
| VDIETLEK | BCO2 | Beta-Carotene Oxygenase 2 | 10 | 25 | 0.87 | 25 | Hs, Bt, Ss, Pt, Pa |
| NETGGQPVLDSPSK | BMPRAI1 | BMP/Retinoic Acid Inducible Neural Specific 1 | 10 | 25 | 0.25 | 25 | Hs, Mm, Rn, Cp, Bt, Ss, Pt, Pa |
| LLPPVSGGYR | CYP26C1 | Cytochrome P450 Family 26 Subfamily C Member 1 | 10 | 25 | 0.77 | 25 | Hs, Mm, Rn, Cp, Bt, Ss,Pt, Dr,XI |
| ENVLITGGGR | DHRS3 | Dehydrogenase/Reductase SDR Family Member 3 | 10 | 25 | 0.76 | 25 | Hs, Cp, Bt, Ss |
| TALLGLTK | DHRS4 | Dehydrogenase/Reductase SDR Family Member 4 | 10 | 25 | 0.92 | 25 | Hs, Mm, Rn, Cp, Bt, Ss, Pt, Pa, XI |
| DELKPFDILQPK | RAI2 | Retinoic Acid Induced 2 | 10 | 10 | 0.62 | 25 | Hs, Pt, Pa |
| GLGQPDLPK | RARG | Retinoic Acid Receptor Gamma | 10 | 25 | 0.82 | 25 | Hs, Mm, Rn, Cp, Bt, Ss, Pt, Pa |
| VGEGFEETVDGR | RBP1 | Retinol Binding Protein 1 | 10 | 25 | 0.79 | 25 | Hs, Mm, Rn, Cp, Bt, Ss, Pt, Pa |
| HEVLGNVGYLR | RBP3 | Retinol Binding Protein 3 | 10 | 10 | 0.70 | 25 | Hs, Mm, Ss, Pt, Pa |
| DPNGLPPEAQK | RBP4 | Retinol Binding Protein 4 | 10 | 25 | 0.86 | 25 | Hs, Pt |
| TVLITGANSGLGR | RDH14 | Retinol Dehydrogenase 14 | 25 | 25 | 0.58 | 25 | Hs, Mm, Rn, Cp, Bt, Ss, Pt, Pa |
| YSPGWDAK | RDH5 | Retinol Dehydrogenase 5 | 10 | 25 | 0.21 | 25 | Hs, Mm,Rn, Cp, Bt, Ss, Pt, Pa, Dr, XI |
| AIHLFPDAK | RXRB | Retinoid X Receptor Beta | 10 | 25 | 0.81 | 25 | Hs, Mm, Rn, Cp, Bt, Ss, Pt, Pa, Dr, XI |
| VTEEEINVLK | ANKRPD1 | Ankyrin Repeat Domain 1 | 50 | 50 | 0.71 | 50 | Hs, Pt |
| GLQVALEEFHK | RARRES2 | Retinoic Acid Receptor Responder 2 | 50 | 50 | 0.83 | 50 | Hs, Cp, Bt, Ss, Pa |
| VAVAASKPHVEIR | RBP1 | Retinol Binding Protein 1 | 10 | 50 | 0.83 | 50 | Hs, Mm,Rn, Cp, Bt, Ss, Pt, Pa, XI |
| QEGDIFYIK | RBP2 | Retinol Binding Protein 2 | 25 | 50 | 0.83 | 50 | Hs, Cp, Bt, Pt, Pa |
| VIDQDGDNFK | RBP2 | Retinol Binding Protein 2 | 10 | 50 | 0.93 | 50 | Hs, Pt, Pa |
| LFALFAR | RDH10 | Retinol Dehydrogenase 10 | 10 | 50 | 0.77 | 50 | Hs, Mm, Rn, Bt, Ss, Pt, Pa, Dr, XI |
| EIQTTTGNQQLVLR | RDH11 | Retinol Dehydrogenase 11 | 10 | 50 | 0.30 | 50 | Hs, Pt, Pa |
| LANVLFTR | RDH12 | Retinol Dehydrogenase 12 | 10 | 10 | 0.67 | 50 | Hs, Rn, Cp, Bt, Ss, Pt, Pa, Dr |
| ELDLASLR | RDH14 | Retinol Dehydrogenase 14 | 10 | 50 | 0.84 | 50 | Hs, Mm, Rn, Cp, Ss |
| YGVFAFSDSLR | RDH16 | Retinol Dehydrogenase 16 | 50 | 50 | 0.89 | 50 | Hs, Mm*, Rn*, Bt, Ss, Pt, Pa |
| TVDSGSLYVR | RDH8 | Retinol Dehydrogenase 8 | 10 | 50 | 0.89 | 50 | Hs, Pt, Pa |
| ATVQSVLLDSAGK | RETSAT | Retinol Saturase | 10 | 50 | 0.98 | 50 | Hs, Cp, Pt, Pa |
| INPTELETIK | RPE65 | Retinoid Isomerohydrolase RPE65 | 25 | 50 | 0.75 | 50 | Hs, Mm, Rn, Cp, Ss, Pt, Pa |
| AIVLFPNPSK | RXRA | Retinoid X Receptor Alpha | 10 | 50 | 0.91 | 50 | Hs, Mm, Rn, Cp,Bt, Ss, Dr, XI |
| TAESGELHGLTDEK | TTR | Transthyretin | 10 | 25 | 0.19 | 50 | Mm, Rn |
| VLAVELASTSR | CYP26B1 | Cytochrome P450 Family 26 Subfamily B Member 1 | 25 | 100 | 0.88 | 100 | Hs, Mm, Rn, Cp, Bt, Ss, Pt, Pa |
| DTHETAAYVR | CYP26C1 | Cytochrome P450 Family 26 Subfamily C Member 1 | 10 | 50 | 0.70 | 100 | Hs, Mm, Rn, Bt, Pt, Pa |
| DAAAGPDVGAAGDAPAPANK | DGAT1 | Diacylglycerol O-Acyltransferase 1 | 25 | 100 | 0.24 | 100 | Hs, Pt, Pa |
| AHAWPSPYK | GPCR5A | G Protein-Coupled Receptor Class C Group 5 Member A | 50 | 100 | 0.88 | 100 | Hs, Pt, Pa |
| YNPESLLQEGEGR | RARRES1 | Retinoic Acid Receptor Responder 1 | 25 | 100 | 0.93 | 100 | Hs, Pa |
| IAVAASKPAVEIK | RBP2 | Retinol Binding Protein 2 | 25 | 50 | 0.19 | 100 | Hs, Mm, Rn, Cp, Bt, Ss, Pt, . XI |
| SENFEEELK | RBP2 | Retinol Binding Protein 2 | 25 | 100 | 0.69 | 100 | Hs, Pt, Pa |
| VVVITGANTGIGK | RDH12 | Retinol Dehydrogenase 12 | 10 | 25 | 0.35 | 100 | Hs, Mm, Rn, Cp, Bt, Ss, Pt, Pa, Dr |
| DLYLPASR | RDH8 | Retinol Dehydrogenase 8 | 25 | 100 | 0.90 | 100 | Hs, Mm, Rn, Bt, Pt, Pa |
| LQYPELFDSLSPFAVR | RLBP1 | Retinaldehyde Binding Protein 1 | 10 | 10 | 0.62 | 100 | Hs, Bt, Pt, Pa |
| IVEAILQEK | SDR16C5 | Short Chain Dehydrogenase/Reductase Family 16C Member 5 | 10 | 50 | 0.30 | 100 | Hs, Pt, Pa |
| ILELIQSGVAEGAK | ALDH1A2 | Aldehyde Dehydrogenase 1 Family Member A2 | 10 | 500 | 0.98 | 500 | Hs, Rn, Cp, Bt, Ss, Pt, Pa |
| FAVPLHYDK | BCO1 | Beta-Carotene Oxygenase 1 | 10 | 25 | 0.24 | 500 | Hs, Mm, Rn, Cp, Ss, Pt, Pa |
| YGFYIK | CYP26A1 | Cytochrome P450 Family 26 Subfamily A Member 1 | 10 | 50 | 0.25 | 500 | Hs, Mm, Rn, Cp, Bt, Ss, Pt, Pa |
| HLEGAISEK | CYP26C1 | Cytochrome P450 Family 26 Subfamily C Member 1 | 50 | 50 | 0.37 | 500 | Hs, Pa |
| VVVITDAISGLGK | DHRS7C | Dehydrogenase/Reductase SDR Family Member 7C | 10 | 500 | 0.86 | 500 | Hs, Mm, Bt, Ss, Pt, Pa, XI |
| LAIVGGGYTPSK | DHRS9 | Dehydrogenase/Reductase SDR Family Member 9 | 500 | 500 | 0.89 | 500 | Hs, Pt, Pa |
| ALLNEEVAR | LRAT | Lecithin Retinol Acyltransferase | 10 | 25 | 0.60 | 500 | Hs, Pt, Pa, Ss |
| TPGPPGLTTTAPPDK | RAI1 | Retinoic Acid Induced 1 | 10 | 25 | 0.30 | 500 | Hs, Pt |
| AGEDPHSFYFPGQFAFSK | RARRES2 | Retinoic Acid Receptor Responder 2 | 10 | 500 | 0.42 | 500 | Hs, Pa |
| ALDIDFATR | RBP2 | Retinol Binding Protein 2 | 10 | 10 | 0.67 | 500 | Hs, Mm, Rn, Bt, Ss, XI |
| FSGLWYAIK | RBP4 | Retinol Binding Protein 4 | 10 | 10 | 0.69 | 500 | Mm, Rn |
| GSQVTTYSVHPGTVQSELVR | RDH11 | Retinol Dehydrogenase 11 | 50 | 50 | 0.39 | 500 | Hs, Pt, Pa |
| IHFHNLQGEK | RDH11 | Retinol Dehydrogenase 11 | 500 | 500 | 0.95 | 500 | Hs, Mm, Cp, Bt, Ss, Pt, Pa |
| HLDLASLK | RDH13 | Retinol Dehydrogenase 13 | 25 | 500 | 0.71 | 500 | Hs, Bt, Pt |
| TVIVTGANTGIGK | RDH13 | Retinol Dehydrogenase 13 | 10 | 10 | 0.18 | 500 | Hs, Mm, Rn, Cp, Bt |
| LANILFTR | RDH14 | Retinol Dehydrogenase 14 | 50 | 500 | 0.92 | 500 | Hs, Mm, Rn, Cp, Bt, Ss, Pt, Pa,Dr, XI |
| FGLEAFSDSLR | RDH5 | Retinol Dehydrogenase 5 | 10 | 500 | 0.79 | 500 | Hs, Mm, Rn, Cp, Bt, Pt, Pa |
| AIVLFPNDAK | RXRG | Retinoid X Receptor Gamma | 100 | 500 | 0.86 | 500 | Hs, Mm, Rn, Cp, Bt, Ss, Pt, Pa, Dr* |
| GLQSSYSEELYR | STRA6 | Stimulated by Retinoic Acid 6 | 50 | 500 | 0.91 | 500 | Hs, Bt, Pt, Pa |
| ATPQLPAQLQLEHR | STRA8 | Stimulated by Retinoic Acid 8 | 10 | 500 | 0.36 | 500 | Hs, Pt |
| AADDTWEPFASGK | TTR | Transthyretin | 10 | 50 | 0.28 | 500 | Hs |
| QAFQISPPWR | ALDH1A1 | Aldehyde Dehydrogenase 1 Family Member A1 | 500 | 500 | 0.61 | 2000 | Hs, Mm*, Rn*, Cp, Bt, Ss, Pt, Pa, |
| EEIFGPVPILK | ALDH1A3 | Aldehyde Dehydrogenase 1 Family Member A3 | 500 | 2000 | 0.95 | 2000 | Hs, Mm, Rn, Bt, Ss, Pt, Pa |
| GDLANTFR | ARID1A | AT-Rich Interaction Domain 1A | 100 | 2000 | 0.93 | 2000 | Hs, Pt, Pa |
| LTSVPTLR | BCO1 | Beta-Carotene Oxygenase 1 | 500 | 2000 | 0.98 | 2000 | Hs, Mm, Rn, Cp, Pt |
| RPNLQQVTENLIK | BMPRAI2 | BMP/Retinoic Acid Inducible Neural Specific 2 | 50 | 50 | 0.67 | 2000 | Hs, Mm, Rn, Cp, Ss, Pt, Pa |
| NSFYETSSFHR | LRAT | Lecithin Retinol Acyltransferase | 25 | 50 | 0.34 | 2000 | Hs, Pt, Pa |
| LAIVLFTK | RDH13 | Retinol Dehydrogenase 13 | 10 | 2000 | 0.88 | 2000 | Hs, Pt, Pa |
| TL SAVGTPLNALGSPYR | RXRG | Retinoid X Receptor Gamma | 500 | 2000 | 0.91 | 2000 | Hs, Mm, Rn, Cp, Bt, Ss, Pt, Pa |

<sup>1</sup> LOD, limit of detection (attomoles), defined as the lowest peptide abundance from the standard curve for which mean(area) > mean(blank) + 3SD(blank), in a matrix of tryptically digested *S. cerevisiae* protein
<sup>2</sup> LOQ, limit of quantitation (attomoles), defined as the lowest peptide abundance where mean(area) > mean(blank) + 10SD(blank) in said matrix
<sup>3</sup> rdotpLOQ, dot product between light and heavy peptides at the LOQ
<sup>4</sup> PLQ, practical limit of quantitation (attomoles). The minimum abundance from calibration curve where rdotp >0.7
<sup>5</sup> Selected species commonly used in preclinical studies that harbor identical tryptic peptides (SwissProt and TrEMBL) according to MaCPepDB (https://macpepdb.mpc.rub.de/). Bt, Bos taurus; Dr, Danio rerio; Hs, Homo sapiens; Mm, Mus musculus; Pongo abelii; Pt, Pan troglodytes; Rn, Rattus norvegicus; Ss, Sus Scrofa; XI, Xenopus laevis

### Linearity, sensitivity and reproducibility of peptide calibration curves

The intensity and spread of the 80 heavy peptides across the 60-min gradient is shown in **Figure 2A**. Calibration curves were generated by spiking light synthetic peptides at 10, 25, 50, 100, 500, and 2000 attomoles in the presence of a fixed 50 femtomoles heavy internal standard. Targeted calibration curves showed linear quantitative performance across the tested dynamic range (10–2000 attomoles). After normalization to heavy internal standards, most peptides displayed strong linear responses, with high coefficients of determination (R^2^) between 0.93 to 1.0. A representative calibration curve and chromatograms for the peptide DLYLPASR (RDH8) is shown in **Figure 2B-C**, and calibration curves for all peptides in **Table 1** are shown in **Supporting Figure S1**. We determined the practical limit of quantitation (PLQ) for all 80 peptides in the assay. Eight peptides achieved a PLQ of 2,000 attomoles, 21 peptides reached 500 attomoles, 11 peptides achieved 100 attomoles, 15 peptides reached 50 attomoles, 14 peptides achieved 25 attomoles, and 11 peptides demonstrated a PLQ of 10 attomoles **(Table 1)**. Analytical precision was evaluated by calculating the coefficient of variation (CV) for each peptide at 2000 attomoles, a concentration above or equal to PLQ for all targets. Among the 80 peptides analyzed, ∼73% exhibited CV values ≤10%, while the remaining ∼27% had CV values between 11% and 21% **(Supporting Figure S2).**

**Figure 2.**
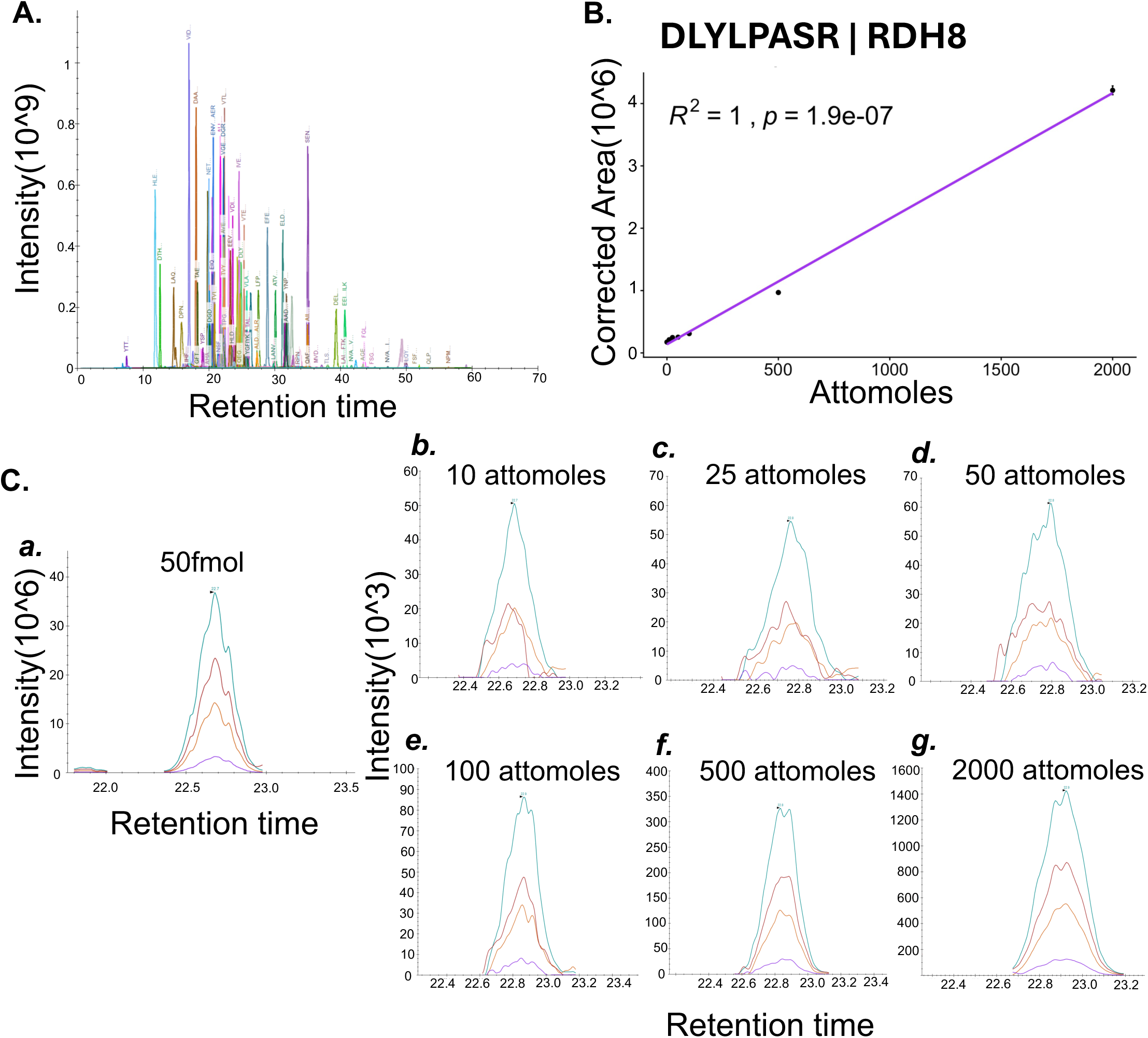
SureQuant targeted acquisition performance. **(A)** Integrated MS¹ extracted ion chromatogram peak areas for all 80 heavy internal standard peptides across the SureQuant method.**(B)** Representative calibration curve for the peptide DLYLPASR (RDH8). Light peptides were spiked at 10, 25, 50, 100, 500, and 2000 attomoles in the presence of a fixed 50 fmol heavy internal standard. The response was linear across the dynamic range (R^2^ = 1.0)**.(C)** Representative chromatograms for DLYLPASR showing **(*a*)** the heavy internal standard peptide and **(*b–g*)** the corresponding light peptide at each calibration level (10–2000 attomoles), demonstrating concentration-dependent increases in signal intensity. Traces show intensity of four product ions from the peptide, color matched across all graphs.

### Targeted quantification of retinoid pathway proteins across human-derived tissues and cell types using the SureQuant method

We employed a diverse panel of human cell types and tissues to investigate retinoid metabolism, including A549 lung carcinoma cells, HepG2 hepatocellular carcinoma cells, retinal organoids (ROs), retinal pigment epithelium (RPE) cells, and healthy adult heart tissue lysates **(Figure 3A)**. Hierarchical clustering of log2-transformed, median-centered, z-scored light/heavy ratios grouped samples according to tissue of origin **(Figure 3B)**, demonstrating that the relative abundances of retinoid pathway proteins vary substantially between tissues and establish distinct local retinoid metabolic networks. Absolute quantification values (fmol/mg protein) for all measured proteins are provided in **Supporting Table S1**.

**Figure 3.**
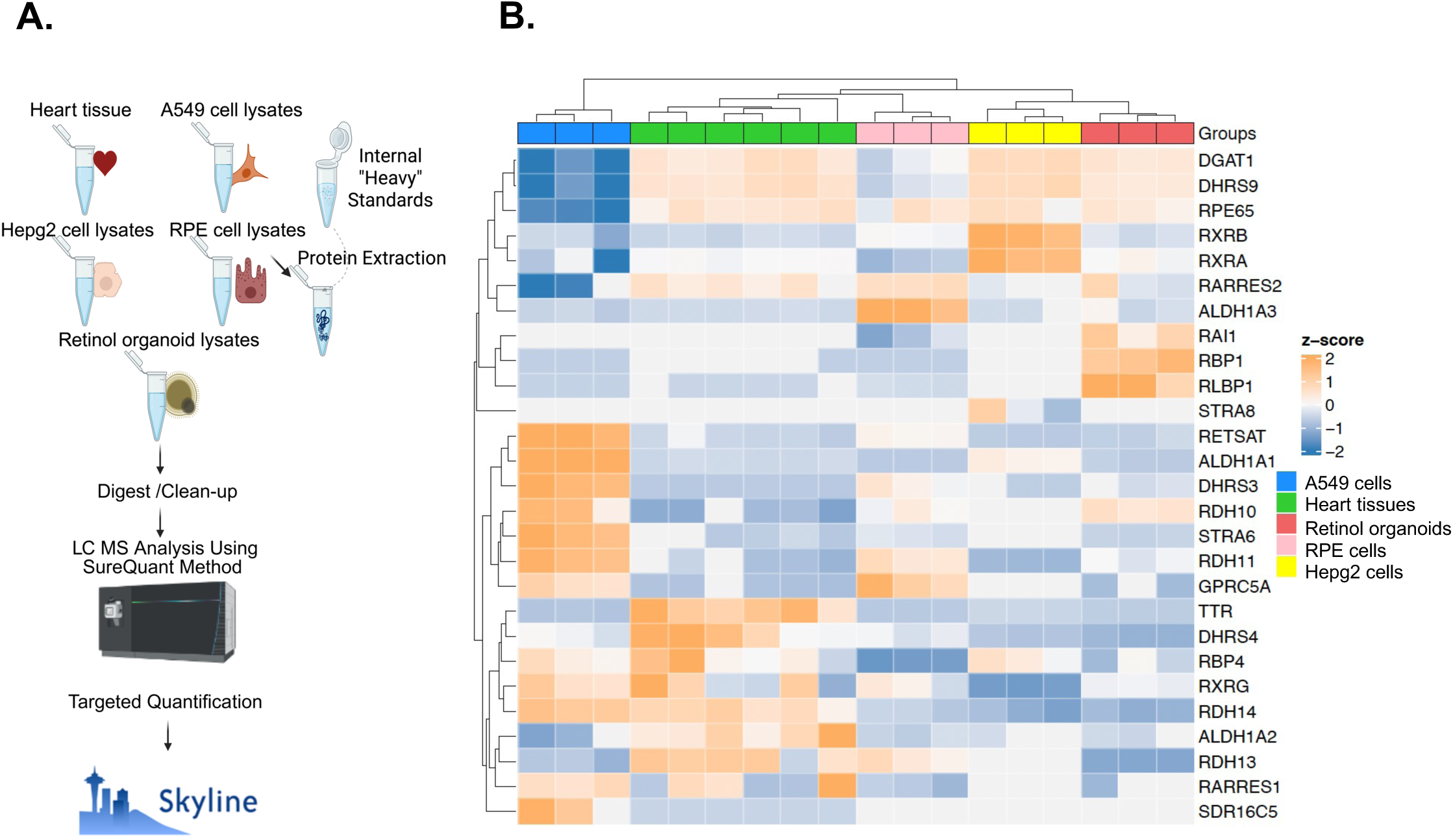
Targeted quantification of retinoid pathway proteins across human-derived tissues and cell types using the SureQuant method. **(A)** Workflow of sample preparation and LC–MS analysis. **(B)** Heat map of *z*-scored light/heavy peptide ratios across samples. Colors represent row-wise relative protein abundance (z-scores), with blue indicating relatively lower and orange indicating relatively higher abundance across samples. Light gray blocks denote missing values corresponding to signals below the limit of detection. For hierarchical clustering, values below detection were imputed as low abundance to stabilize clustering, while missing values are displayed as gray in the heat map and do not indicate absolute absence or presence.

Starting from the beginning of the retinoid signaling pathway, STRA6 showed its highest abundance in A549 cells, consistent with enhanced retinol uptake capacity in this cell type. RBP4 exhibited the highest abundance in A549 cells, followed by adult heart tissue and HepG2 cells, with lower levels observed in ROs. TTR was most abundant in adult heart tissue, likely reflecting the high vascularity and circulating protein content of the heart. Next, intracellular retinoid-binding proteins showed distinct tissue-specific distributions. RBP1 was relatively higher in retinal organoids, while RBP2 was relatively higher in RPE cells. SDR16C5, RDH10, RDH11, and RDH14 showed the highest abundance in A549 cells, followed by adult heart tissue, whereas RDH13 was highest in heart tissue. DHRS3 showed the highest abundance in A549 cells, while DHRS4 was abundant in both A549 and heart tissues. ALDH1A1 was highly abundant in A549 cells, ALDH1A2 in heart tissue, and ALDH1A3 in RPE cells. Looking at RXRs, RXRA and RXRB showed the highest abundance in HepG2 and A549 cells, RXRG was most abundant in A549 cells. RLBP1 and RPE65, as expected, were highest in RO and RPE cells, respectively. Regarding retinoid-responsive proteins, STRA8 was detected only in HepG2 cells. GPRC5A and RARRES2 showed the highest abundance in RPE cells, whereas RARRES1 was highest in A549 cells **(Figure 4)**.

**Figure 4.**
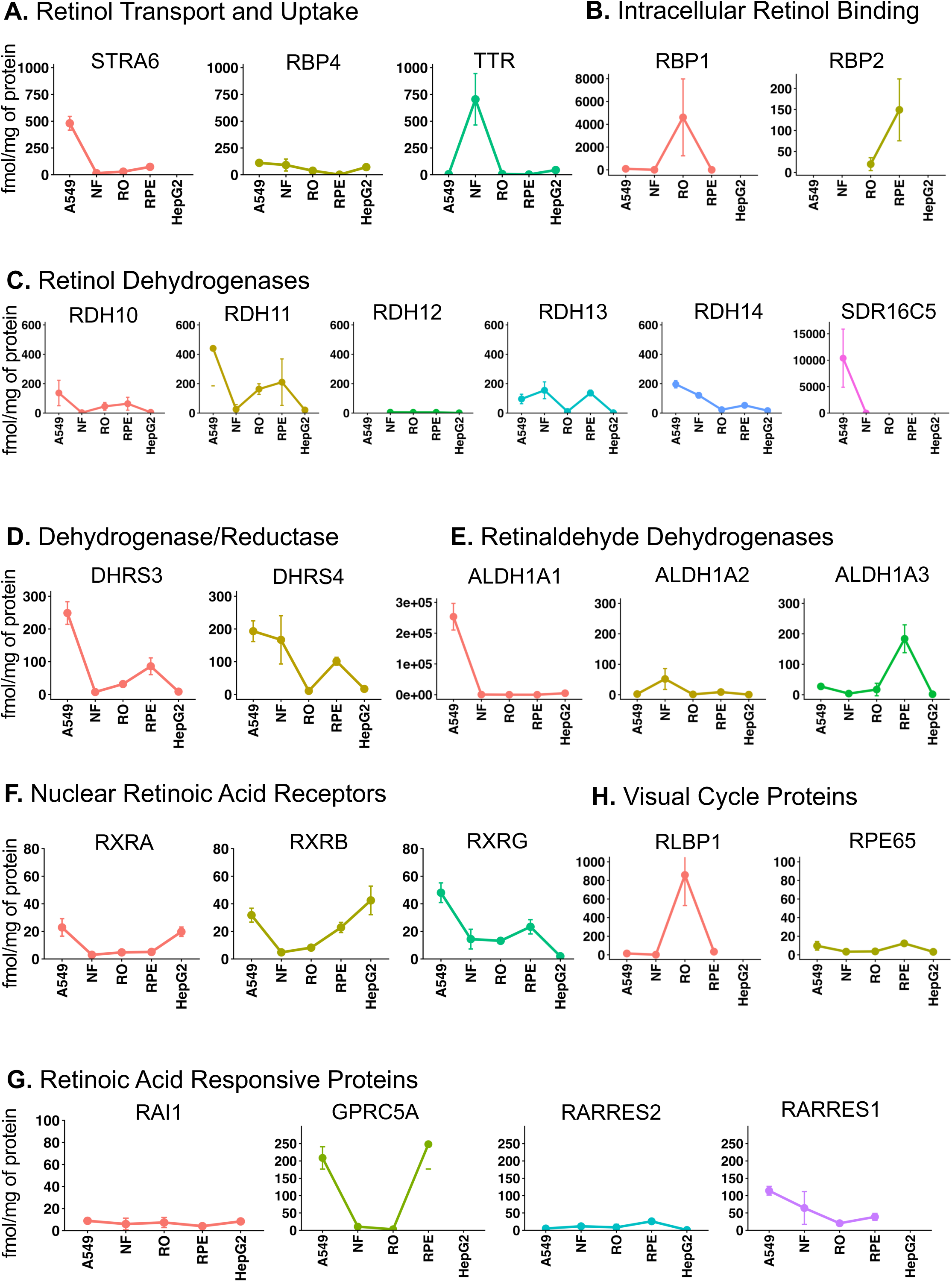
Functional category–based quantification of retinoid pathway proteins across cell types. Line graphs showing the abundance of peptides corresponding to proteins grouped by functional category across A549 cells, adult non-failing heart tissue (NF), retinal organoids (RO), retinal pigment epithelium (RPE), and HepG2 cells. Values represent direct peptide abundance (fmol/mg of total protein) measurements without normalization or scaling. Replicate (n=3 per group) measurements within each cell type were averaged, and error bars represent standard deviation.

### Abundance of retinoid pathway proteins in mouse hearts during embryonic, postnatal, adult development

Given the important roles of retinoid signaling in cardiac development and its later transition toward regulating adaptive and stress-responsive pathways, we investigated the expression dynamics of retinoid signaling proteins across three stages of mouse cardiac development: embryonic, postnatal, and adult. At embryonic day 9.5 (E9.5), heart tubes were dissected together with the adjacent posterior region, where ALDH1A2—a key RA-synthesizing enzyme—is robustly expressed during early cardiogenesis (34). These samples, together with postnatal day 1 (P1) hearts and adult (12-week) hearts, were processed for SureQuant-targeted acquisition of retinoid pathway proteins **(Figure 5-A)**. Principal component analysis (PCA) of proteins quantified demonstrated clear separation among embryonic, postnatal, and adult cardiac samples, highlighting stage-specific remodeling of the retinol metabolism network during cardiac maturation **(Figure 5-B)**. Of the 41 peptides monitored by this method in Mus Musculus species, 22 are shown in the heatmap. Hierarchical clustering of log^2^ transformed, median centered, z-scored abundances revealed distinct expression patterns switches in retinoid metabolism proteins across the three developmental time points, with clear separation between embryonic, postnatal, and adult stages **(Figure 5C).**

**Figure 5.**
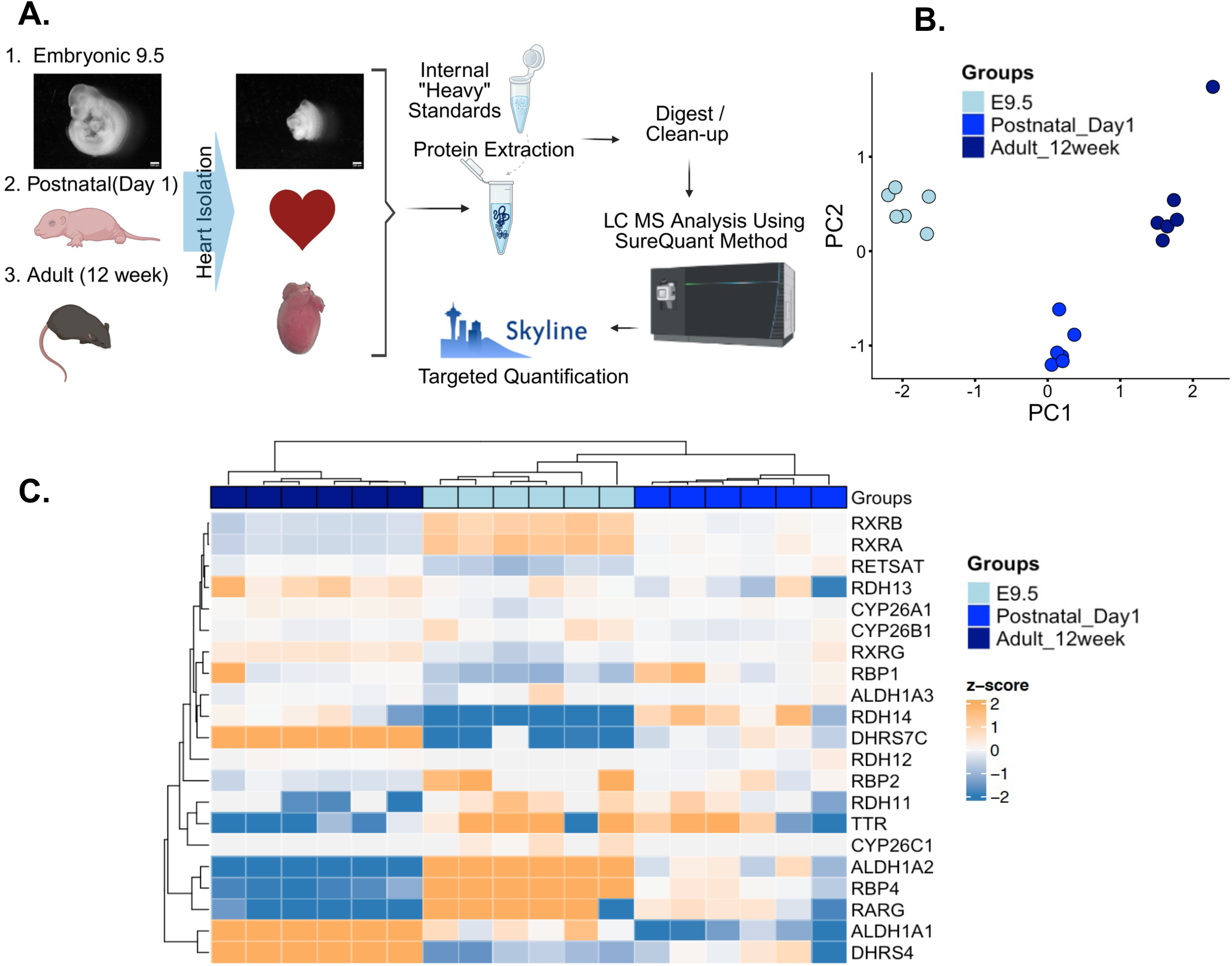
Targeted quantification of retinoid pathway proteins in mouse hearts across three developmental time points. **(A)** Workflow of sample preparation and LC–MS analysis. Mouse hearts were collected at three time points: embryonic day 9.5 (E9.5), postnatal day 1 (P1), and adult (12 weeks). For each time point, *n* = 6 hearts were processed. **(B)** Unsupervised PCA biplot of all samples **(C)** Heat map of light/heavy peptide ratios across samples. Colors represent row-wise relative protein abundance (z-scores), with blue indicating relatively lower and orange indicating relatively higher abundance across samples. Light gray blocks denote missing values corresponding to signals below the limit of detection. For hierarchical clustering, values below detection were imputed as low abundance to stabilize clustering, while missing values are displayed as gray in the heat map and do not indicate absolute absence or presence.

Starting from the upstream retinol signaling pathway, both RBP4 and TTR were highest during the embryonic stage and gradually decreased after birth. Among intracellular retinol-binding proteins, RBP1 was also highest at the embryonic stage. RBP2 was detected only during embryonic development. Several developmental changes were observed among the RDHs. RDH10 and RDH11 were relatively higher during embryonic development and decreased postnatally and into adulthood, whereas RDH14 increased in postnatal and adult hearts. Within the SDR family, DHRS4 and DHRS7C became more abundant in adult hearts. Among the ALDH enzymes, ALDH1A2 was relatively higher at E9.5 and decreased in postnatal and adult hearts, while ALDH1A1 showed the opposite trend and increased during later developmental stages., RXRA, RXRB, and RARG were highest during the embryonic stage, whereas RXRG was relatively higher in adult hearts. CYP26A and CYP26B were also more abundant during embryonic development **(Figure 6)**. Taken together, these expression dynamics underscore a highly active retinoid signaling axis during embryogenesis, highlighting its temporal regulation during cardiac development. The raw abundances (fmol/mg of protein) for all proteins are present in **Supporting Table S2**. To orthogonally validate the targeted proteomics findings, Western blot analyses was performed DHRS4 in mouse hearts at E9.5, P1, and 12 weeks of age. The developmental changes observed were consistent with those measured by SureQuant PRM **(Supporting Figure S3)**

**Figure 6.**
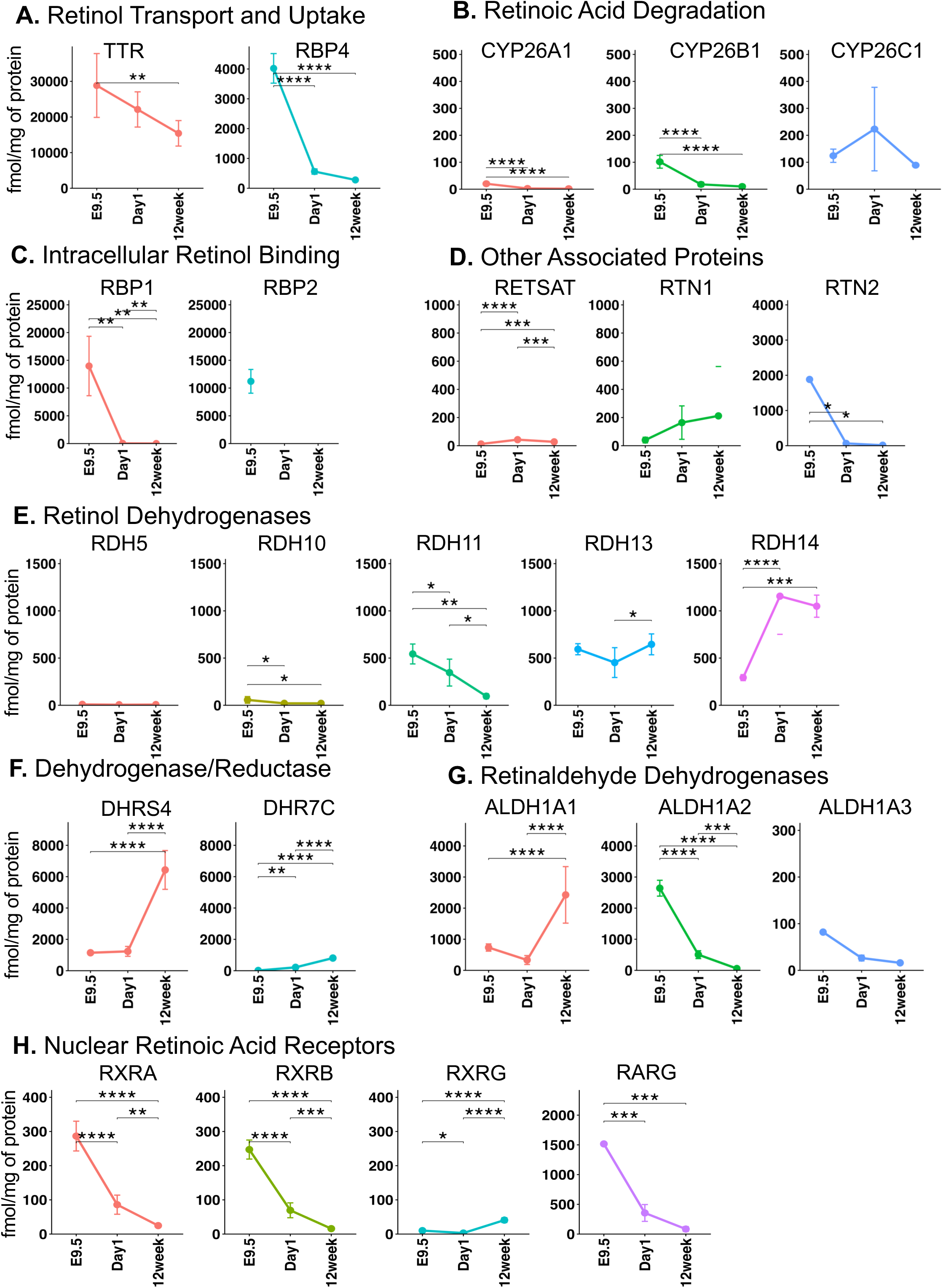
Functional category–based quantification of retinoid pathway proteins across developmental mouse heart tissues. Line graphs showing the abundance of peptides corresponding to proteins grouped by functional category across embryonic mouse hearts (E9.5), postnatal day 1 hearts, and adult mouse hearts at 12 weeks. Values represent direct peptide abundance measurements (fmol/mg of total protein) obtained without normalization or scaling. Replicate measurements (n=6 per group) within each developmental group were averaged, and error bars represent standard deviation. Statistical significance was assessed using one-way ANOVA followed by pairwise post hoc comparisons. Significance is indicated as follows: * p < 0.05, ** p < 0.01, *** p < 0.001, and **** p < 0.0001.

### Abundance of retinoid pathway proteins in rat hearts during postnatal day 1 and day 14

Finally, we explored the cross-species applicability of the SureQuant assay by applying the panel to rat heart tissue collected during early postnatal development (postnatal days 1 and 14) **(Figure 7A).** During postnatal cardiac development, RDH14 was the most abundant detected member of the RDH family, followed by RDH13, whereas RDH8, RDH10, and RDH5 were detected at substantially lower abundances. Among the ALDH family proteins, ALDH1A1 exhibited the highest abundance, followed by ALDH1A2 and ALDH1A3 **(Supporting Table S3).** Interestingly, ALDH1A1 abundance increased significantly from postnatal day 1 to day 14, whereas ALDH1A2 abundance decreased over the same developmental period. Western blot analysis confirmed a significant reduction in ALDH1A2 protein expression between postnatal day 1 and day 14 (p = 0.01), consistent with the decrease detected by the SureQuant assay (p = 0.004). In addition, RXRG expression increased significantly during this developmental interval **(Figure 7B-C)**. Together, these findings indicate dynamic remodeling of retinoid metabolism and signaling pathways during early postnatal cardiac maturation. Western blot analysis of DHRS4 in rat pup hearts at postnatal days 1 and 14, presented in **Supporting Figure S4**, further validated the corresponding SureQuant findings.

**Figure 7.**
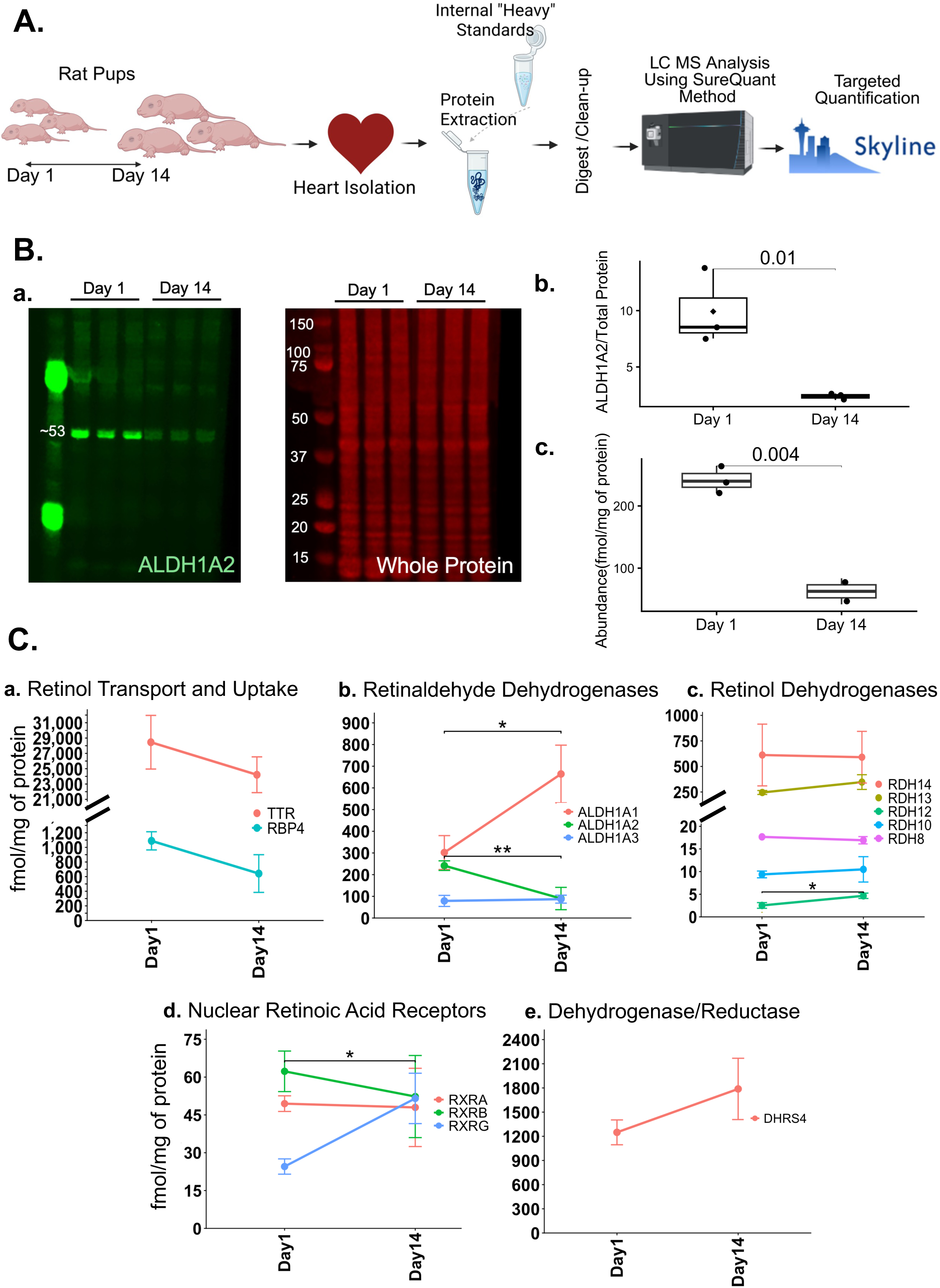
Targeted quantification of retinoid pathway proteins across Rat pup heart at day 1 and day 14 using the SureQuant method. **(A)** Overview of the sample preparation and LC–MS/MS workflow. **(B)** Representative Western blot showing **ALDH1A2** band at approximately 53 kDa. (a) shows a representative Western blot, panel (b) shows protein abundance determined by Western blotting and normalized to total protein staining. (c) shows protein abundance measured by SureQuant PRM and reported as fmol/mg of total protein. Data are presented as mean ± SD. Statistical significance between P1 and P14 groups was assessed using an unpaired two-sided t-test. Significant differences are indicated on the plots (p < 0.05). **(C)** Peptide abundances corresponding to retinoid pathway proteins grouped by functional category in Day 1 and Day 14 postnatal rat hearts. Each developmental stage included three biological replicates (n = 3), except for RDH8 at Day 1, where only one replicate yielded a detectable signal. Values represent absolute peptide abundances (fmol/mg total protein) obtained directly by SureQuant analysis without normalization or scaling. Data are presented as mean ± SD. Statistical significance between developmental stages was assessed using two-sided t-tests. Significance is indicated as follows: * p < 0.05, ** p < 0.01, *** p < 0.001, and **** p < 0.0001.

## Discussion

### Practical aspects of SureQuant assay

SureQuant™ IS-PRM provides highly sensitive and accurate peptide quantification at attomole–femtomole levels without the need for target enrichment. This approach is dynamic and robust to retention time variability compared with scheduled PRM. When the distinctive fragment ions for a corresponding synthetic internal standard (the trigger peptide) are detected, the mass spectrometer gathers MS/MS scans for the target peptide (35). The constant switch between watch mode and quant mode allows the instrument to monitor for trigger peptides while simultaneously acquiring quantitative data only when targets are present. In this enhanced acquisition process, the watch mode analyzes both MS1 and MS2 data to track the elution of the internal standard(22). Because SureQuant continuously searches for the trigger peptide rather than relying on predefined retention time windows, the method is inherently robust to retention time shifts over time. As a result, retention time calibration is not required, and changes in chromatographic gradients or column replacement can be performed without concern for missing peptide elution windows, as may occur with scheduled PRM methods. These features provide improved duty cycle efficiency while reducing the risk of target peptides drifting outside scheduled acquisition windows. Notably, all parameters necessary for the workflow are established in a single survey run and can be applied to all subsequent SureQuant analyses of the same peptide panel, thereby streamlining assay development and implementation(33). Internal standard peptides serve a dual function as limit-of-detection controls; detection of only the heavy internal standard without the corresponding light peptide indicates the endogenous peptide is either absent or present below quantifiable levels(33).

SureQuant has been successfully applied in diverse contexts, including high-throughput biomarker monitoring in body fluids(22) validation and quantification of peptide antigens presented on MHC molecules(36), and targeted analysis of signaling networks such as tyrosine phosphorylation pathways(37), particularly in settings where detection of low-abundance proteins remains challenging using conventional approaches.

### Tissue-specific organization of retinoid metabolism and its developmental regulation in the heart

Prior transcriptomic studies and large-scale expression resources such as the Human Protein Atlas have shown substantial tissue-specific variation in genes encoding retinoid metabolic enzymes and binding proteins(38–40). Despite this knowledge, systematic characterization of these proteins has proven difficult, as conventional bottom-up proteomics workflows frequently fail to detect or accurately quantify many components of the retinoid pathway. We employed a diverse panel of human cell types and tissues to investigate retinoid metabolism, including A549 lung carcinoma cells, HepG2 hepatocellular carcinoma cells, retinal organoids (ROs), retinal pigment epithelium (RPE) cells, and healthy adult heart tissue lysates. These models were selected to represent the diverse physiological functions of retinoid signaling across different organ systems. In the liver, vitamin A is converted to all-trans retinoic acid (ATRA) to regulate fatty acid metabolism and bile acid production and is also stored as retinyl esters(41–44). Photoreceptor cells depend on the vitamin A derivative 11-cis-retinal to capture light and initiate phototransduction, while the retinal pigment epithelium (RPE) continuously regenerates 11-cis-retinal from all-trans retinol through the enzymatic reactions of the visual cycle, ensuring sustained visual function(45). In the heart, retinoid signaling is critical for maintaining cardiac homeostasis, with dysregulation linked to pathological remodeling and heart failure(3).

Proteins such as STRA6, RBP4, RXRG, and several RDH family members (SDR16C5, RDH10, RDH11, and RDH14) exhibited their highest abundance in A549 cells **(Figure 4)**. The visual cycle proteins RLBP1 and RPE65 were predominantly enriched in retinal organoids and RPE cells, respectively, consistent with their established roles in retinal retinoid recycling **(Figure 4)**. Differential enrichment was also observed among retinaldehyde dehydrogenases, with ALDH1A1 highest in A549 cells, ALDH1A2 enriched in heart tissue, and ALDH1A3 most abundant in RPE cells, suggesting tissue-specific routes of retinoic acid synthesis. Additional tissue-dependent differences were observed across dehydrogenase/reductase (DHRS) enzymes, nuclear retinoic acid receptor isoforms, and retinoid-responsive proteins. DHRS3 was most abundant in A549 cells, while DHRS4 was enriched in both A549 cells and heart tissues. RXRA and RXRB showed the highest abundance in A549 and HepG2 cells **(Figure 4)**.

RA signaling plays a critical role at multiple stages of cardiac development, including anterior–posterior patterning of the cardiac mesoderm, specification of cardiomyocyte lineages, epicardial formation, outflow tract morphogenesis, expansion of the ventricular compact myocardium, and coronary vessel development(34). Postnatally, however, RA signaling appears to transition from regulating developmental morphogenesis(46) to supporting maintained functions like regulation of metabolic flexibility through PDK4/PDH pathways(47). Our data further support this concept by providing evidence of tissue specificity and a cardiac developmental switch across multiple steps of the retinoid metabolic pathway—from retinol to retinal to all-trans retinoic acid (ATRA), as well as its downstream nuclear signaling and degradation. Beginning with the first enzymatic step, we examined the retinol dehydrogenase (RDH) family, which catalyzes the conversion of retinol to retinal in the initial stage of retinoic acid biosynthesis **(Figure 8).**

**Figure 8.**
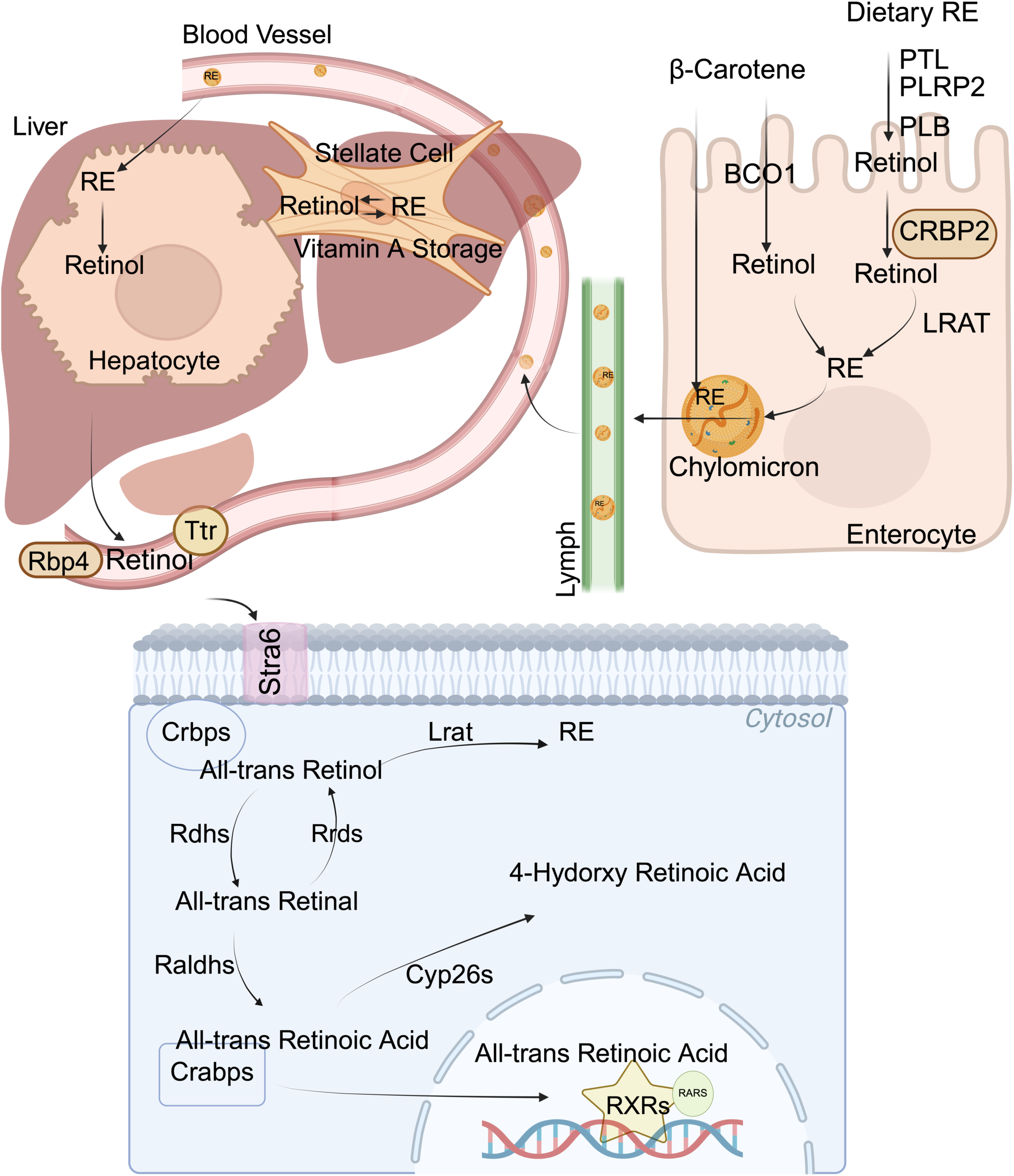
Vitamin A Absorption, Storage, Signaling and Metabolism. Retinyl esters (RE) are hydrolyzed to retinol in the intestinal lumen by pancreatic (PTL, PLRP2) and brush border (PLB) enzymes. Retinol enters enterocytes via passive diffusion, binds to CRBP2, and is re-esterified by lecithin retinol acyltransferase (LRAT). RE are then packaged into chylomicrons with other dietary lipids and apoB. Provitamin A carotenoids (e.g., β-carotene) are absorbed by enterocytes and either converted to retinol by BCO1 and retinal reductase or remain intact. Both RE and carotenoids are secreted into the lymph via chylomicrons and subsequently enter the bloodstream. Approximately 70% of chylomicron-bound retinyl esters are taken up by the liver for storage. Once inside hepatocytes, retinyl esters are hydrolyzed back to all-trans retinol, which can then enter hepatic stellate cells for storage. Alternatively, all-trans retinol can be released into circulation. Circulating retinol bound to retinol-binding protein 4 (RBP4) and transthyretin (TTR) is transported into cells through the membrane receptor stimulated by retinoic acid 6 (STRA6). In the cytosol, retinol associates with cellular retinol-binding proteins (CRBPs) and can be stored as retinyl esters through the activity of LRAT. Retinol is oxidized to retinal by retinol dehydrogenases (RDHs), while members of the short-chain dehydrogenase/reductase (SDR) family (DHRS) catalyze the reverse reaction, reducing retinal back to retinol. Retinal is irreversibly converted to all-trans retinoic acid (ATRA) by retinaldehyde dehydrogenases (RALDHs), encoded by ALDH1A genes. Intracellular ATRA can bind cellular retinoic acid-binding proteins (CRABPs) and either be degraded by cytochrome P450 family 26 enzymes (CYP26) or translocate to the nucleus, where it regulates gene transcription through retinoic acid receptors (RARs) and retinoid X receptors (RXRs).

Examining RDH abundance across developmental stages revealed notable shifts in the relative abundance of specific isoforms. RDH10 enzymes play a critical role during embryogenesis(48) with RDH10 knockout embryos dying around E10.5 with severe RA deficiency phenotypes affecting cardiac and other organ development(48–50). Our data indicate that RDH10 had significantly higher abundances in embryonic period and RDH14 becomes more abundant in the postnatal and adult mouse heart **(Figure 6)**, although the exact function of RDH14 in adult heart retinoid metabolism has not been fully explored in previous studies. Counteracting the enzymatic activity of RDHs, members of the DHRS family (also known as retinal reductases or RRDs) catalyze the reverse reaction, reducing retinal back to retinol (51) **(Figure 8)**. When examined across mouse cardiac developmental stages, DHRS4 showed highest abundance observed in adult mouse hearts, suggesting a potential role in maintaining retinoid homeostasis in mature cardiac tissue **(Figure 6)**. Consistent with this observation, DHRS4 abundance increased by approximately 1.5-fold between postnatal day 1 and day 14 in rat hearts, indicating that this developmental increase begins during the early postnatal period **(Figure 7).**

Downstream in the retinoid pathway, retinal is irreversibly oxidized to ATRA by members of the retinaldehyde dehydrogenase (RALDH) family, encoded by ALDH1A genes(52). During cardiac development, ALDH1A2 is clearly the dominant isoform during embryogenesis, consistent with its well-established role in cardiac patterning and morphogenesis(53). In our study, analysis of whole-heart lysates from embryonic, postnatal, and adult mouse hearts revealed a shift from high ALDH1A2 abundance during embryonic development to predominance of ALDH1A1 in the adult heart**(Figure 6)**. To further define the timing of this transition, we examined rat hearts during the early postnatal period (postnatal days 1 and 14). Interestingly, ALDH1A1 abundance increased while ALDH1A2 abundance decreased between day 1 and day 14, suggesting that this developmental switch occurs within the first two weeks after birth **(Figure 7)**. Once synthesized, retinoic acid can either be degraded by CYP26 enzymes, (54), or translocate to the nucleus where it activates transcription through retinoic acid receptors (RARs) and retinoid X receptors (RXRs)(55) **(Figure 8)**. In our study, in mouse hearts, both CYP26A and CYP26B appeared to be relatively more abundant during embryonic stages. Regarding nuclear receptors, RXRA and RXRB show higher abundance during embryonic stages in mouse hearts, with expression decreasing after birth while RXRG had higher abundance in adult mouse hearts.

Together, these observations suggest a coordinated remodeling of the retinoid metabolic and signaling network across cardiac development. The shift in enzyme abundance from embryonic to postnatal and adult stages may reflect a transition in retinoid metabolism from developmental signaling toward roles that support metabolic and transcriptional regulation in the mature heart, where retinoid signaling has been implicated in regulating antioxidant defenses, calcium handling, metabolic pathways, and hypertrophic remodeling in cardiac tissue(56). Although our study evaluated protein abundance rather than enzymatic activity, and abundance alone does not necessarily reflect functional dominance, the observed developmental patterns suggest a reorganization of the retinoid metabolic network accompanying cardiac maturation.

### Limitations of the using SureQuant Assay in this study

A known limitation of targeted MS approaches, including SureQuant, is the assumption that the spiked stable isotope–labeled standards are recovered and detected consistently across assays. Isotopically labeled peptides do not always behave exactly like the peptide from endogenous proteins, which can lead to small differences in measured light-to-heavy ratios. These effects may introduce minor assay-level bias, especially in complex tissue samples like heart. To account for this, we applied replicate-level normalization by scaling the light-to-heavy ratios in each run according to the median heavy peptide signal of the assay relative to the global median heavy signal across all assays. Furthermore, targeted MS workflows are typically developed around a selected precursor charge state identified during assay optimization or survey scans, with the assumption that this ion species consistently represents the peptide during subsequent analyses. However, peptide ionization behavior and charge-state distributions can vary across sample matrices and acquisition conditions, potentially resulting in incomplete sampling of the total peptide population and underestimation of true abundance. Finally, an inherent limitation of bottom-up proteomics is that proteins are quantified indirectly through surrogate tryptic peptides, and not all peptides exhibit favorable analytical properties for sensitive and reproducible detection(57). Successful targeted peptide assays therefore depend on identifying highly ionizable, high-yield peptides with robust MS performance, which is not always feasible for every protein of interest. In such cases, even carefully optimized assays may rely on peptides that perform sub optimally, which can limit sensitivity or quantitative precision despite consistent sample preparation and instrument settings. For instance, for several proteins of broad interest to the retinoid community, (RARA, RARB), we have, thus far, been unable to identify peptides that match our criteria for high-performance and sensitive protein quantification (e.g. full-scan intensity > 10^7^ at 50 fmol).

## Conclusion

Taken together, we developed a SureQuant parallel reaction monitoring (PRM) method to comprehensively detect and quantify peptides representing enzymes of the retinoic acid metabolism and signaling pathway. The peptide panel is applicable across multiple species, with most peptides validated in human, mouse and rat derived samples. We applied our panel to diverse tissue types and cardiac samples across developmental stages, revealing distinct tissue-specific expression patterns for retinoid metabolic proteins and identifying a developmental switch in retinoid metabolism within the heart—transitioning from roles in cardiac development to functions in adult homeostasis. It is noteworthy that the principles and workflow established here are equally applicable to next-generation targeted quantitation methods, including hybrid PRM/DIA approaches(58), offering a scalable framework for future investigations of retinoid signaling networks in health and disease.

## Supporting Information and Data Availability

This article contains supporting information including figures and tables. Skyline documents and associated raw mass spectrometry in this study have been deposited in Panorama Public and are publicly available at https://panoramaweb.org/Foster_Retinoid_Surequant.url

