## Supporting Information for "SureQuant™ IS-PRM Enables Cross-Species Targeted Quantification of Retinoid Metabolism and Signaling Proteins in the Heart"

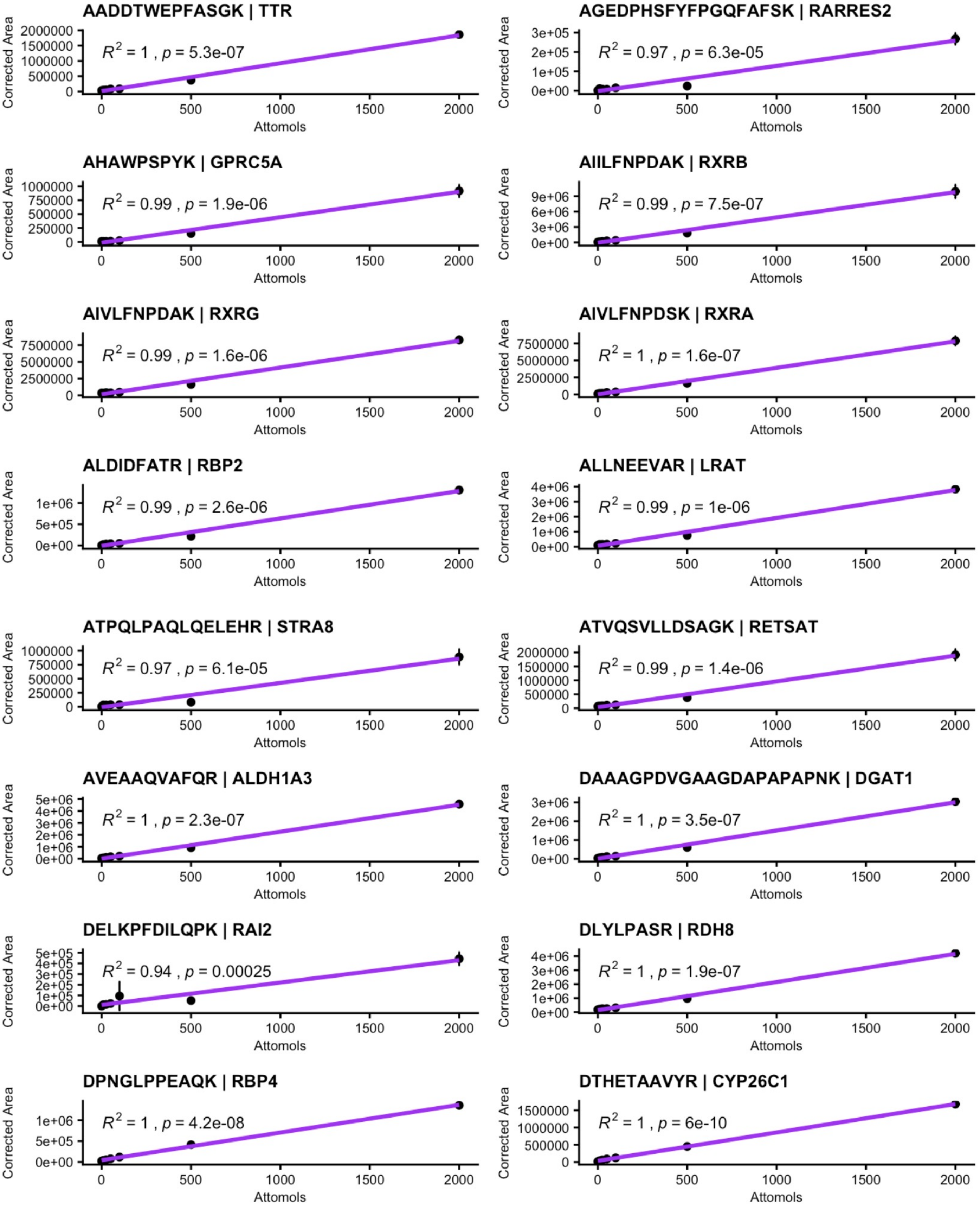
AADDTWEPFASGK | TTR
R2 = 1 , p = 5.3e-07
AGEDPHSFYFPGQFAFSK | RARRES2
R2 = 0.97 , p = 6.3e-05
AHAWPSPYK | GPRC5A
R2 = 0.99 , p = 1.9e-06
AIILFNPDAK | RXRB
R2 = 0.99 , p = 7.5e-07
AIVLFNPDAK | RXRG
R2 = 0.99 , p = 1.6e-06
AIVLFNPDSK | RXRA
R2 = 1 , p = 1.6e-07
ALDIDFATR | RBP2
R2 = 0.99 , p = 2.6e-06
ALLNEEVAR | LRAT
R2 = 0.99 , p = 1e-06
ATPQLPAQLQELEHR | STRA8
R2 = 0.97 , p = 6.1e-05
ATVQSVLLDSAGK | RETSAT
R2 = 0.99 , p = 1.4e-06
AVEAAQVAFQR | ALDH1A3
R2 = 1 , p = 2.3e-07
DAAAGPDVGAAGDAPAPAPNK | DGAT1
R2 = 1 , p = 3.5e-07
DELKPFDILQPK | RAI2
R2 = 0.94 , p = 0.00025
DLYLPASR | RDH8
R2 = 1 , p = 1.9e-07
DPNGLPPEAQK | RBP4
R2 = 1 , p = 4.2e-08
DTHETAAVYR | CYP26C1
R2 = 1 , p = 6e-10
Corrected Area
Attomols

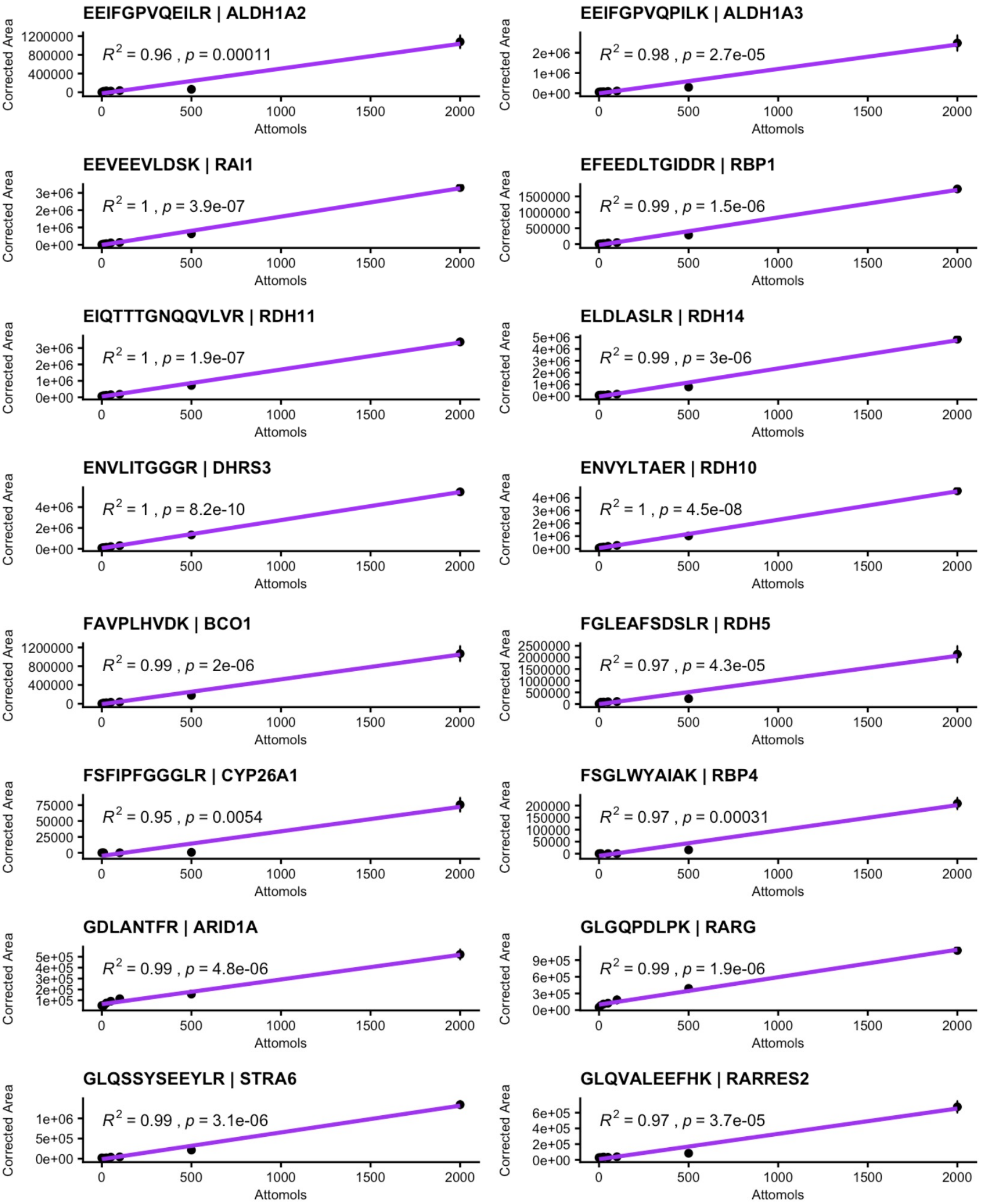

EEIFGPVQEILR | ALDH1A2
R2 = 0.96 , p = 0.00011
EEIFGPVQPILK | ALDH1A3
R2 = 0.98 , p = 2.7e-05
EEVEEVLDSK | RAI1
R2 = 1 , p = 3.9e-07
EFEEDLTGIDDR | RBP1
R2 = 0.99 , p = 1.5e-06
EIQTTTGNQQVLVR | RDH11
R2 = 1 , p = 1.9e-07
ELDLASLR | RDH14
R2 = 0.99 , p = 3e-06
ENVLITGGGR | DHRS3
R2 = 1 , p = 8.2e-10
ENVYLTAER | RDH10
R2 = 1 , p = 4.5e-08
FAVPLHVDK | BCO1
R2 = 0.99 , p = 2e-06
FGLEAFSDSLR | RDH5
R2 = 0.97 , p = 4.3e-05
FSFIPFGGGLR | CYP26A1
R2 = 0.95 , p = 0.0054
FSGLWYAIAK | RBP4
R2 = 0.97 , p = 0.00031
GDLANTFR | ARID1A
R2 = 0.99 , p = 4.8e-06
GLGQPDLPK | RARG
R2 = 0.99 , p = 1.9e-06
GLQSSYSEEYLR | STRA6
R2 = 0.99 , p = 3.1e-06
GLQVALEEFHK | RARRES2
R2 = 0.97 , p = 3.7e-05
Corrected Area
Attomols
0
500
1000
1500
2000

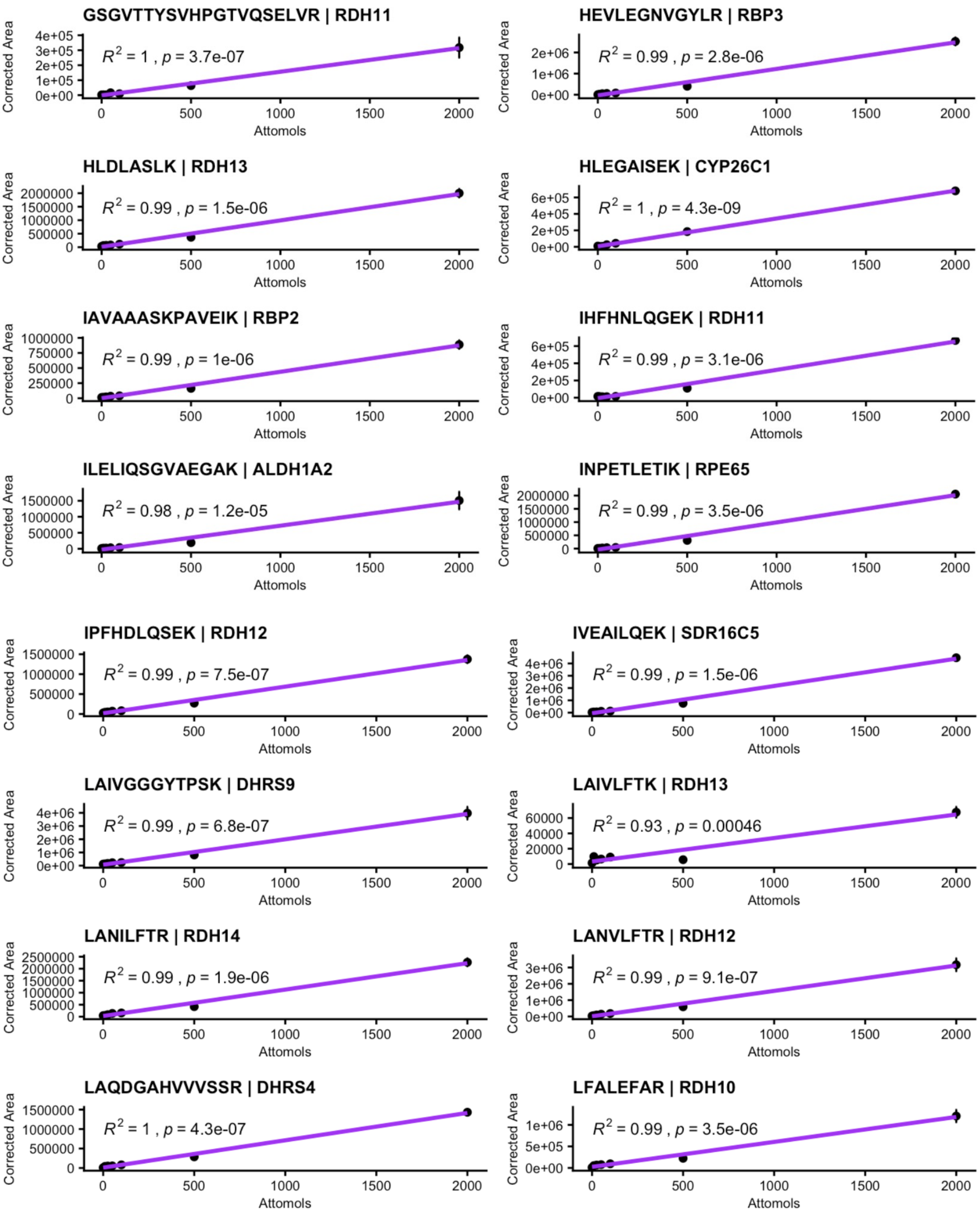
GSGVTTYSVHPGTVQSELVR | RDH11
$R^2$ = 1 , $p$ = 3.7e-07
HEVLEGNVGYLR | RBP3
$R^2$ = 0.99 , $p$ = 2.8e-06
HLDLASLK | RDH13
$R^2$ = 0.99 , $p$ = 1.5e-06
HLEGAISEK | CYP26C1
$R^2$ = 1 , $p$ = 4.3e-09
IAVAAASKPAVEIK | RBP2
$R^2$ = 0.99 , $p$ = 1e-06
IHFHNLQGEK | RDH11
$R^2$ = 0.99 , $p$ = 3.1e-06
ILELIQSGVAEGAK | ALDH1A2
$R^2$ = 0.98 , $p$ = 1.2e-05
INPETLETIK | RPE65
$R^2$ = 0.99 , $p$ = 3.5e-06
IPFHDLQSEK | RDH12
$R^2$ = 0.99 , $p$ = 7.5e-07
IVEAILQEK | SDR16C5
$R^2$ = 0.99 , $p$ = 1.5e-06
LAIVGGGYTPSK | DHRS9
$R^2$ = 0.99 , $p$ = 6.8e-07
LAIVLFTK | RDH13
$R^2$ = 0.93 , $p$ = 0.00046
LANILFTR | RDH14
$R^2$ = 0.99 , $p$ = 1.9e-06
LANVLFTR | RDH12
$R^2$ = 0.99 , $p$ = 9.1e-07
LAQDGAHVVVSSR | DHRS4
$R^2$ = 1 , $p$ = 4.3e-07
LFALEFAR | RDH10
$R^2$ = 0.99 , $p$ = 3.5e-06
Corrected Area
Attomols

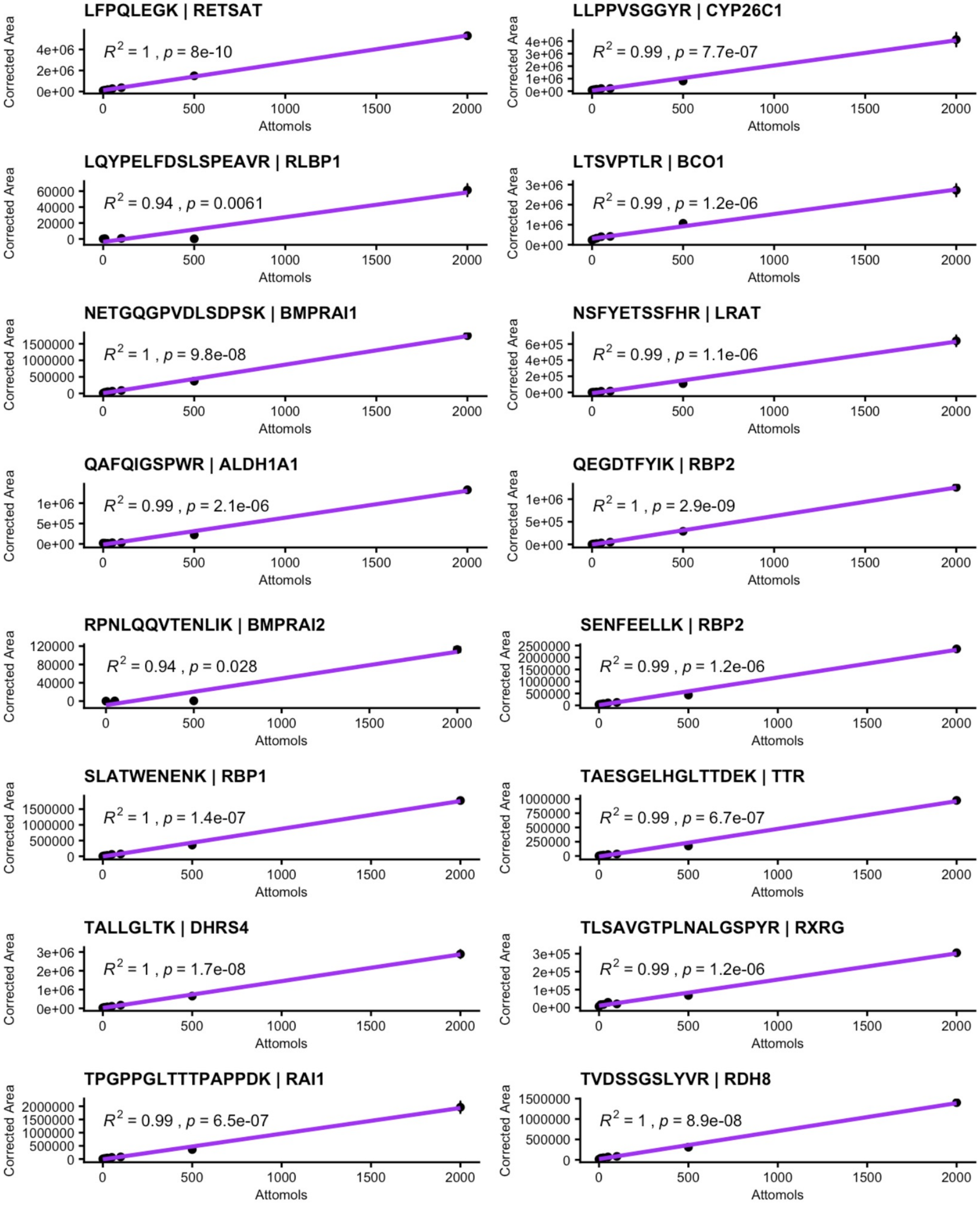
LFPQLEGK | RETSAT
$R^2 = 1$ , $p$ = 8e-10
LLPPVSGGYR | CYP26C1
$R^2 = 0.99$ , $p$ = 7.7e-07
LQYPELFDSLSPEAVR | RLBP1
$R^2 = 0.94$ , $p$ = 0.0061
LTSVPTLR | BCO1
$R^2 = 0.99$ , $p$ = 1.2e-06
NETGQGPVDLSDPSK | BMPRAI1
$R^2 = 1$ , $p$ = 9.8e-08
NSFYETSSFHR | LRAT
$R^2 = 0.99$ , $p$ = 1.1e-06
QAFQIGSPWR | ALDH1A1
$R^2 = 0.99$ , $p$ = 2.1e-06
QEGDTFYIK | RBP2
$R^2 = 1$ , $p$ = 2.9e-09
RPNLQQVTENLIK | BMPRAI2
$R^2 = 0.94$ , $p$ = 0.028
SENFEELLK | RBP2
$R^2 = 0.99$ , $p$ = 1.2e-06
SLATWENENK | RBP1
$R^2 = 1$ , $p$ = 1.4e-07
TAESGELHGLTTDEK | TTR
$R^2 = 0.99$ , $p$ = 6.7e-07
TALLGLTK | DHRS4
$R^2 = 1$ , $p$ = 1.7e-08
TLSAVGTPLNALGSPYR | RXRG
$R^2 = 0.99$ , $p$ = 1.2e-06
TPGPPGLTTTPAPPDK | RAI1
$R^2 = 0.99$ , $p$ = 6.5e-07
TVDSSGSLYVR | RDH8
$R^2 = 1$ , $p$ = 8.9e-08
Corrected Area
Attomols
0
500
1000
1500
2000

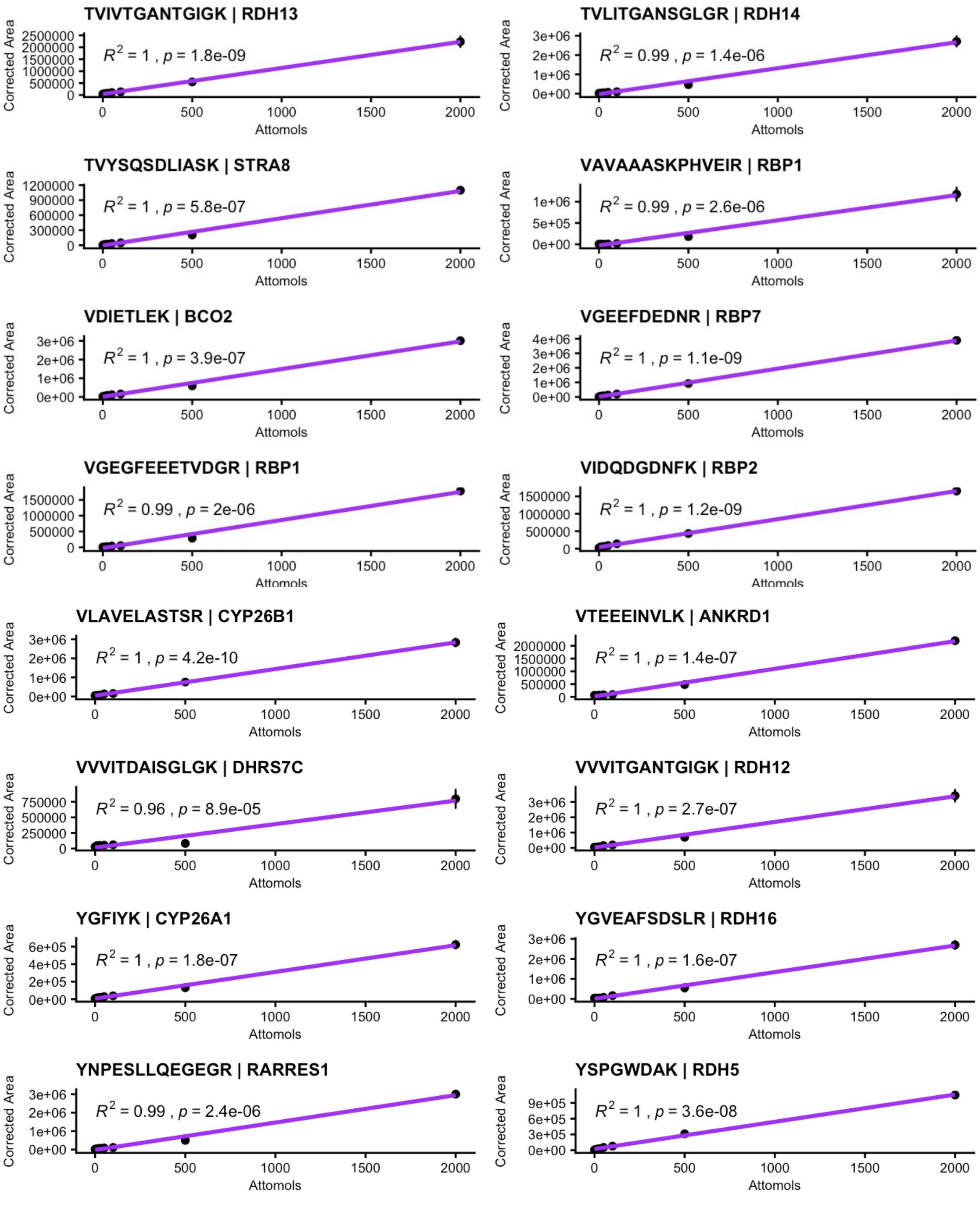


**Supporting Figure S1 Calibration curves for all peptides generated using the SureQuant workflow.** Light peptide standards were prepared at concentrations of 0, 10, 25, 50, 100, 500, and 2000 attomoles per vial, with a constant 50 fmol of heavy peptide added as an internal standard. The x-axis indicates the amount of light peptide (attomoles), and the y-axis shows the measured peak area ratio of light to heavy peptide. Data points represent mean values from technical replicates, with error bars indicating standard deviation.

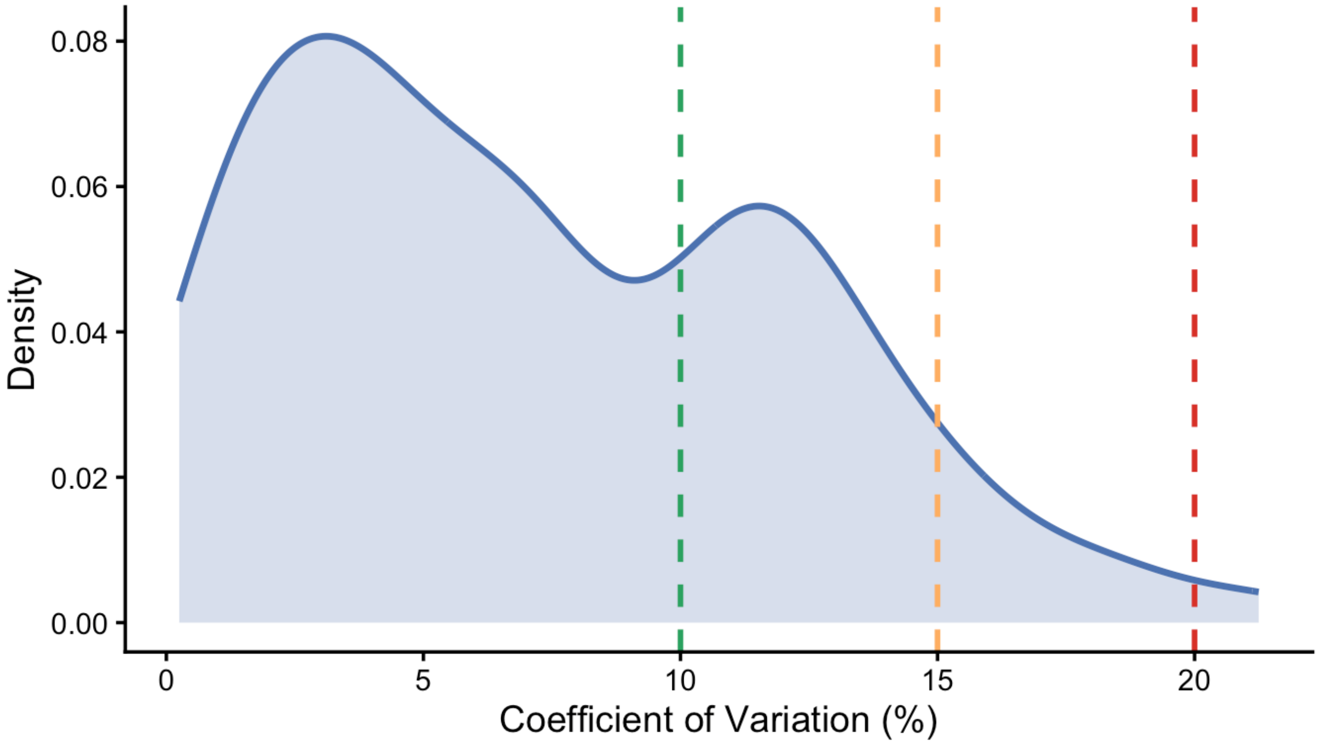


**Supporting Figure S2. Density distribution of the coefficient of variation (CV%).** CV was calculated for each peptide at 2000 attomoles of light peptide spiked into a constant background of 50 fmol heavy internal standard, using three replicate injections, and derived from the resulting median light-to-heavy ratios. Among the 80 peptides analyzed, 72.5% exhibited CV values ≤10%, indicating high quantitative reproducibility, while the remaining 27.5% showed CV values between 11% and 21%. Dashed vertical lines indicate reproducibility thresholds (10%, 15%, and 20% CV). The majority of peptides fall within the 0-10 % CV range, demonstrating robust quantitative performance of the assay at concentrations above or equal to the practical limit of quantification (PLQ).

**Peptide Quantification (fmol/mg of protein)**

Values scaled for visualization

| protein | peptide | A549_1 | A549_2 | A549_3 | NF_1 | NF_2 | NF_3 | NF_4 | NF_5 | NF_6 | RO1 | RO2 | RO3 | RPE1 | RPE2 | RPE3 | Hepg2_1 | Hepg2_2 | Hepg2_3 |
|---|---|---|---|---|---|---|---|---|---|---|---|---|---|---|---|---|---|---|---|
| ALDH1A1 | QAFQIGSPWR | 255,893.20 | 208,840.85 | 295,389.64 | 478.13 | 486.66 | 421.29 | 683.72 | 637.62 | 604.87 | 245.13 | 20.27 | 8.75 | 10.48 | 9.46 | 13.71 | 4,001.98 | 6,374.47 | 4,294.54 |
| RBP1 | SLATWENENK | 20.96 | 29.58 | 25.64 | NA | NA | NA | NA | 1.36 | NA | 5,375.93 | 6,426.18 | 10,136.94 | 1.63 | 6.16 | 2.39 | NA | NA | NA |
| SDR16C5 | IVEAILQEK | 6,495.73 | NA | 14,275.44 | 6.43 | 5.67 | 10.89 | 9.35 | 16.22 | 10.57 | NA | NA | NA | NA | NA | NA | NA | NA | NA |
| RBP1 | VGEGFEEETVDGR | 29.84 | 421.84 | 26.51 | NA | 4.70 | NA | 11.04 | NA | NA | 1,578.28 | 1,686.62 | 2,442.74 | NA | NA | NA | NA | NA | NA |
| TTR | AADDTWEPFASGK | 6.61 | 4.81 | 4.97 | 713.50 | 927.88 | 1,021.68 | 639.12 | 372.43 | 552.86 | 9.52 | 5.93 | 5.30 | 1.96 | 2.98 | 2.83 | 36.77 | 49.28 | 46.62 |
| RDH11 | EIQTTTGNQQVLVR | 636.15 | 620.83 | 749.71 | 44.02 | 44.89 | NA | 105.78 | 12.35 | NA | 216.74 | 151.49 | 200.35 | 320.07 | 411.86 | 321.89 | 15.66 | 24.82 | 21.51 |
| RLBP1 | LQYPELFDSLSPEAVR | 6.67 | 17.56 | 19.10 | 0.43 | 2.48 | NA | 0.88 | NA | 1.66 | 1,032.69 | 1,063.82 | 479.51 | 33.15 | 45.00 | 23.18 | NA | NA | NA |
| DHRS4 | TALLGLTK | 184.94 | 166.67 | 228.24 | 157.19 | 76.54 | 259.39 | 169.98 | 97.17 | 240.81 | 11.00 | 13.44 | 7.67 | 95.45 | 115.37 | 93.37 | 14.91 | 19.28 | 16.33 |
| RDH13 | TVIVTGANTGIGK | 119.65 | 109.52 | 141.42 | 161.23 | 62.27 | 164.17 | 274.05 | 172.06 | 183.77 | 12.15 | 5.42 | 7.56 | 139.65 | 159.33 | 136.63 | 1.79 | 2.87 | 1.78 |
| STRA6 | GLQSSYSEEYLR | 456.59 | 432.16 | 553.59 | 12.26 | 40.23 | NA | NA | 1.16 | 5.79 | 28.46 | 25.44 | 32.67 | 74.64 | 89.52 | 57.62 | NA | NA | NA |
| RETSAT | ATVQSVLLDSAGK | 341.92 | 321.92 | 405.30 | 22.02 | 23.38 | 29.02 | 73.60 | 12.18 | 29.26 | 16.64 | 17.43 | 33.85 | 98.58 | 109.79 | 97.45 | 4.99 | 10.86 | 7.33 |
| RDH14 | LANILFTR | 199.80 | 200.31 | 239.76 | 113.15 | 109.38 | 145.55 | 130.78 | 115.35 | 155.21 | 25.67 | 20.59 | 18.15 | 47.73 | 58.20 | 43.87 | NA | NA | NA |
| RDH14 | ELDLASLR | 167.18 | 178.91 | 189.66 | 101.57 | 113.44 | 122.75 | 123.05 | 84.44 | 136.01 | 29.31 | 19.60 | 23.38 | 57.46 | 62.31 | 46.18 | 4.80 | 29.47 | 15.13 |
| RDH13 | LAIVLFTK | 70.01 | 59.13 | 75.31 | 163.07 | 51.83 | 172.93 | 159.64 | 124.52 | 167.62 | 12.88 | 12.18 | 12.76 | 123.47 | 154.92 | 106.15 | NA | NA | NA |
| GPRC5A | AHAWPSPYK | 187.41 | 192.90 | 245.77 | 9.75 | 7.40 | 13.68 | 14.55 | 7.07 | NA | 4.83 | NA | 0.86 | 227.28 | 327.66 | 189.51 | NA | NA | NA |
| RDH11 | GSGVTTYSVHPGTVQSELVR | 210.54 | 174.61 | 250.53 | 4.09 | 10.48 | 9.92 | 12.25 | 3.84 | 7.68 | 138.48 | 130.21 | 141.87 | 65.23 | 82.35 | 59.57 | NA | NA | NA |
| RBP4 | DPNGLPPEAQK | 98.55 | 102.14 | 132.36 | 57.06 | 71.10 | 134.44 | 182.32 | 34.89 | 66.92 | 14.53 | 64.24 | 34.32 | 1.20 | 0.98 | 1.25 | 52.74 | 87.33 | 74.07 |
| RDH10 | LFALEFAR | 194.69 | 208.72 | 239.67 | 1.90 | 1.68 | 3.45 | 3.47 | 1.78 | 2.81 | 68.58 | 66.05 | 74.07 | 102.49 | 124.73 | 75.83 | NA | 6.06 | 3.75 |
| DHRS3 | ENVLITGGGR | 226.71 | 230.99 | 288.53 | 7.52 | 5.27 | 8.13 | 6.52 | 6.90 | 9.29 | NA | 27.94 | 35.48 | 83.49 | 112.93 | 61.71 | 7.64 | NA | 9.38 |
| RARRES1 | YNPESLLQEGEGR | 100.85 | 125.62 | 116.04 | 23.26 | 25.06 | 30.70 | 85.53 | 142.98 | 77.77 | 20.21 | NA | NA | 40.28 | 47.90 | 28.11 | NA | NA | NA |
| RBP2 | QEGDTFYIK | NA | NA | NA | NA | NA | NA | NA | NA | NA | 44.27 | 21.53 | 30.21 | 200.05 | 254.74 | 178.52 | NA | NA | NA |
| ALDH1A3 | EEIFGPVQPILK | 26.29 | 25.44 | 30.19 | 2.61 | 4.58 | 3.76 | 4.77 | 2.72 | 4.46 | 40.26 | 0.94 | 10.30 | 178.31 | 232.06 | 140.98 | NA | 2.50 | 0.87 |
| DHRS4 | LAQDGAHVVVSSR | 52.27 | 56.31 | 64.70 | 54.11 | 28.95 | 87.14 | 93.24 | 30.11 | 77.34 | 4.34 | 2.80 | 2.57 | 29.88 | 43.60 | 34.13 | 5.95 | 8.53 | 6.33 |
| RXRG | AIVLFNPDAK | 39.83 | 51.70 | 52.59 | 7.72 | 17.05 | 25.19 | 18.54 | 6.66 | 11.26 | 12.38 | 13.90 | 13.21 | 23.12 | 28.62 | 18.28 | 1.45 | 2.55 | 1.91 |
| ALDH1A2 | ILELIQSGVAEGAK | 2.44 | NA | 2.13 | 23.66 | 47.62 | 31.14 | 33.42 | 117.00 | 58.19 | 1.58 | 0.90 | NA | 3.41 | 11.16 | 11.96 | NA | 0.48 | NA |
| RXRB | AIILFNPDAK | 28.85 | 28.88 | 37.56 | 4.39 | 3.94 | 4.91 | 5.79 | 5.04 | 4.69 | 9.28 | 6.70 | 8.63 | 21.97 | 26.87 | 19.67 | 33.57 | 53.83 | 40.01 |
| RDH10 | ENVYLTAER | 56.63 | 52.91 | 66.70 | 2.39 | 2.78 | 2.38 | 2.46 | 0.60 | NA | 22.08 | 20.83 | 21.87 | 28.05 | 27.07 | 23.16 | NA | NA | NA |
| RBP2 | IAVAAASKPAVEIK | NA | NA | NA | NA | NA | NA | NA | NA | NA | 12.23 | 2.91 | 7.85 | 83.89 | 112.93 | 65.39 | NA | NA | NA |
| RXRA | AIVLFNPDSK | 25.90 | 15.57 | 27.12 | 1.38 | 1.25 | 3.64 | 4.35 | 3.76 | 3.59 | 3.73 | 6.61 | 3.97 | 4.72 | 5.99 | 4.56 | 17.18 | 23.63 | 18.33 |
| RARRES2 | AGEDPHSFYFPGQFAFSK | 4.70 | NA | 5.71 | 7.16 | 11.58 | 14.85 | 10.78 | NA | 13.56 | 18.52 | 4.14 | 2.63 | 24.56 | 28.64 | 24.44 | NA | 0.87 | NA |
| CYP26B1 | VLAVELASTSR | NA | NA | NA | NA | NA | NA | NA | NA | NA | 4.07 | 43.59 | 74.16 | 0.25 | 0.64 | NA | NA | NA | NA |
| RAI1 | TPGPPGLTTTPAPPDK | 7.16 | 10.21 | 9.39 | 15.54 | 2.21 | 1.94 | 2.84 | 4.87 | 8.61 | 2.85 | 11.94 | 7.45 | 3.88 | 4.81 | 3.55 | 6.05 | 9.49 | 9.62 |
| RPE65 | INPETLETIK | 5.12 | 9.65 | 14.36 | 1.63 | 2.04 | 1.23 | 5.63 | 5.61 | 4.01 | 4.33 | 4.45 | 2.27 | 15.11 | 10.35 | 11.44 | NA | 3.68 | 2.70 |
| RDH12 | VVVITGANTGIGK | NA | NA | NA | 9.01 | 17.21 | 1.86 | 1.28 | 1.93 | 2.06 | 1.98 | 4.26 | 7.98 | 3.97 | 3.61 | 6.49 | 0.76 | 2.05 | 1.33 |
| RDH12 | LANVLFTR | NA | NA | NA | NA | NA | NA | NA | NA | NA | 2.90 | 27.01 | 32.66 | 0.93 | NA | 1.96 | NA | NA | NA |
| RAI1 | EEVEEVLDSK | NA | NA | NA | NA | NA | NA | NA | NA | NA | 10.95 | 5.81 | 10.09 | 6.95 | 8.15 | 8.78 | NA | NA | NA |
| CYP26A1 | FSFIPFGGGLR | NA | NA | NA | NA | NA | NA | NA | NA | NA | 10.53 | 12.24 | NA | 25.25 | NA | NA | NA | NA | NA |
| DGAT1 | DAAAGPDVGAAGDAPAPAPNK | 2.28 | 5.92 | 2.30 | 2.31 | 2.42 | 2.75 | 0.69 | 1.06 | 1.91 | 2.65 | 2.34 | 2.72 | 3.99 | 2.63 | 4.70 | 2.12 | 3.14 | 2.02 |
| BMPRAI1 | NETGQGPVDLSDPSK | NA | 1.23 | 2.50 | 2.99 | 3.12 | 3.66 | 1.29 | 1.26 | 3.86 | 2.33 | 1.83 | 2.44 | 6.04 | 4.85 | 3.38 | 1.83 | 2.30 | 2.37 |
| RBP7 | VGEEFDEDNR | NA | NA | NA | 1.71 | 3.29 | 3.75 | 2.95 | 1.23 | 1.74 | NA | NA | NA | 2.37 | 3.17 | 3.79 | 2.94 | 4.57 | 4.45 |
| RARRES2 | GLQVALEEFHK | 3.77 | 1.09 | 3.11 | 2.10 | 3.47 | 3.46 | 4.73 | NA | 1.28 | 2.73 | 1.69 | 0.96 | 2.56 | 2.89 | 1.79 | NA | NA | NA |
| RBP2 | ALDIDFATR | NA | NA | NA | NA | NA | NA | NA | NA | NA | 0.47 | 1.38 | 0.89 | 5.64 | 9.67 | 4.99 | NA | 7.53 | 4.68 |
| DHRS7C | VVVITDAISGLGK | NA | NA | 0.61 | 1.58 | 3.84 | 12.29 | 3.70 | 6.75 | 2.38 | NA | NA | NA | NA | NA | NA | NA | 2.76 | 1.07 |
| BCO1 | FAVPLHVDK | 4.49 | 4.64 | 7.43 | NA | 3.00 | 1.36 | 0.94 | 1.04 | 1.58 | 1.69 | NA | 0.98 | NA | 0.80 | 2.40 | NA | NA | NA |
| DHRS9 | LAIVGGGYTPSK | 1.32 | 2.19 | 1.59 | 1.62 | 2.55 | 2.22 | 1.49 | 2.24 | 1.93 | 0.97 | 1.11 | 1.49 | 0.76 | 1.15 | 0.80 | 2.73 | 2.08 | 1.73 |
| RAI2 | DELKPFDILQPK | 12.93 | NA | NA | 3.93 | NA | 5.10 | NA | 2.69 | 4.30 | NA | NA | NA | NA | NA | NA | NA | NA | NA |
| RDH12 | IPFHDLQSEK | NA | NA | NA | NA | NA | NA | NA | NA | NA | 1.92 | 6.45 | 10.88 | 1.29 | NA | 1.02 | 0.64 | 1.83 | 1.39 |
| RDH8 | DLYLPASR | 4.33 | NA | 5.80 | 0.62 | 1.16 | 1.81 | NA | NA | 1.71 | NA | NA | NA | NA | NA | NA | NA | NA | NA |
| RBP3 | HEVLEGNVGYLR | NA | NA | NA | NA | NA | NA | NA | NA | NA | 1.25 | 6.14 | 4.79 | 0.98 | NA | NA | NA | NA | NA |
| CYP26C1 | LLPPVSGGYR | NA | NA | NA | NA | NA | NA | NA | NA | NA | 0.28 | NA | NA | 3.20 | 4.63 | 3.52 | NA | NA | NA |
| RDH5 | FGLEAFSDSLR | 3.64 | 4.14 | 0.57 | 0.50 | NA | 0.98 | 0.86 | NA | 0.78 | NA | NA | NA | NA | NA | NA | NA | NA | NA |
| STRA8 | TVYSQSDLIASK | NA | NA | NA | NA | NA | NA | NA | NA | NA | NA | NA | NA | NA | NA | NA | 1.89 | 3.27 | 2.80 |
| RDH8 | TVDSSGSLYVR | NA | 5.27 | NA | 0.66 | 0.38 | NA | NA | NA | NA | NA | NA | NA | NA | NA | NA | NA | NA | 0.98 |
| RDH16 | YGVEAFSDSLR | NA | NA | NA | NA | NA | NA | NA | NA | NA | NA | 1.90 | 0.23 | 1.46 | 1.96 | 1.35 | NA | NA | NA |

**Supporting Table S1 : Table of raw protein abundances (fmol/mg protein) across various human-derived tissues and cell types .** Light-to-heavy ratios were calculated using the three most intense fragment ion transitions for each peptide and determined as Σlight/Σheavy. Endogenous peptide abundances were calculated by scaling these ratios to the known heavy peptide input (50 fmol per 5 µg total protein) and are reported as fmol per mg of total protein. Data are shown for A549 cells, retinal organoids (RO), retinal pigment epithelium (RPE) cells, HepG2 cells (n = 3 biological replicates per group), and non-failing human heart tissue lysates (NF; n = 6 biological replicates). Values represent unnormalized peptide abundances obtained by SureQuant targeted proteomics analysis.

| Peptide Quantification (fmol/mg of protein)<br>Values scaled for visualization | | | | | | | | | | | | | | | | | | | |
|---|---|---|---|---|---|---|---|---|---|---|---|---|---|---|---|---|---|---|---|
| protein | peptide | E95_1 | E95_2 | E95_3 | E95_4 | E95_5 | E95_6 | P1 | P2 | P3 | P4 | P5 | P6 | M1 | M2 | M3 | M4 | M5 | M6 |
| TTR | TAESGELHGLTTDEK | 30,440.47 | 34,664.08 | 39,851.18 | 31,029.44 | 22,129.30 | 14,933.16 | 17,924.29 | 23,087.28 | 14,637.52 | 25,771.70 | 27,817.24 | 23,465.66 | 16,806.84 | 12,337.85 | 19,846.30 | 18,618.87 | 10,876.20 | 14,007.34 |
| RBP1 | VAVAAASKPHVEIR | 17,496.51 | 20,725.00 | 12,386.98 | 17,250.30 | 8,463.54 | 7,522.78 | 33.75 | 27.84 | 19.37 | 48.75 | 32.34 | 40.00 | NA | 4.67 | 4.13 | 7.97 | NA | 7.11 |
| RBP2 | IAVAAASKPAVEIK | 12,395.40 | 14,498.00 | 10,213.97 | 11,997.80 | 9,228.84 | 8,939.60 | NA | NA | NA | NA | NA | NA | NA | 32.88 | NA | NA | NA | NA |
| DHRS4 | TALLGLTK | 1,094.05 | 1,282.04 | 1,312.21 | 1,035.98 | 1,022.72 | 1,125.54 | 1,527.23 | 1,422.33 | 643.01 | 1,348.01 | 1,299.91 | 1,161.42 | 5,710.68 | 5,169.29 | 6,081.46 | 6,607.84 | 8,749.32 | 6,260.92 |
| RBP4 | FSGLWYAIAK | 3,864.97 | 4,793.19 | 4,372.23 | 3,920.28 | 3,806.50 | 3,365.58 | 564.31 | 562.23 | 396.97 | 635.59 | 642.44 | 560.69 | 293.27 | 246.02 | 357.87 | 276.25 | 296.45 | 203.72 |
| ALDH1A1 | QAFQIGSPWR | 618.27 | 686.13 | 836.88 | 603.29 | 770.26 | 886.58 | 418.88 | 511.19 | 96.38 | 311.60 | 384.44 | 268.24 | 1,982.34 | 2,020.25 | 1,936.63 | 2,155.59 | 4,269.06 | 2,193.24 |
| ALDH1A2 | EEIFGPVQEILR | 2,607.08 | 2,873.44 | 2,983.78 | 2,369.42 | 2,361.92 | 2,649.30 | 658.85 | 421.29 | 318.41 | 575.67 | 587.89 | 464.06 | 48.65 | 73.53 | 56.32 | 74.97 | 46.57 | 44.66 |
| RDH14 | TVLITGANSGLGR | 293.77 | 295.78 | 348.68 | 264.34 | 259.90 | 295.15 | 1,438.61 | 960.33 | 428.25 | 1,509.71 | 1,261.77 | 1,332.95 | 1,038.39 | 889.89 | 992.17 | 1,247.74 | 1,065.26 | 1,063.57 |
| RARG | GLGQPDLPK | 79.35 | 1,206.74 | 1,566.50 | 2,100.12 | 2,048.55 | 2,095.91 | 325.49 | 421.46 | 81.34 | 459.84 | 430.50 | 410.81 | 69.41 | 68.14 | 48.58 | 81.55 | 165.73 | 71.67 |
| RDH13 | TVIVTGANTGIGK | 544.14 | 700.44 | 621.64 | 558.54 | 571.32 | 565.82 | 659.23 | 380.51 | 192.34 | 544.48 | 474.47 | 462.08 | 565.73 | 531.13 | 597.10 | 701.43 | 835.56 | 641.23 |
| DHRS7C | VVVITDAISGLGK | 18.31 | 3.52 | NA | 2.70 | 16.01 | 14.92 | 266.34 | 286.60 | 111.85 | 230.66 | 209.66 | 173.51 | 693.36 | 712.59 | 764.08 | 983.42 | 757.32 | 929.85 |
| RTN2 | ALDIDFATR | 4,744.55 | NA | NA | 613.22 | 288.59 | NA | 31.85 | 137.87 | 28.49 | 57.44 | 80.98 | 50.91 | 13.90 | 26.89 | 7.99 | 9.00 | 6.26 | 16.68 |
| RDH11 | IHFHNLQGEK | 497.67 | 514.52 | 698.19 | 463.13 | NA | NA | NA | 308.80 | 126.98 | 509.75 | 409.34 | 375.78 | NA | NA | 65.13 | 108.99 | NA | 114.10 |
| CYP26C1 | LLPPVSGGYR | 91.01 | 129.18 | 156.49 | 115.19 | 106.75 | 146.07 | 496.13 | 127.54 | 144.20 | 77.98 | 309.58 | 182.94 | 281.61 | 19.36 | 25.49 | 39.87 | 129.43 | 38.50 |
| RXRA | AIVLFNPDSK | 262.31 | 302.57 | 366.43 | 243.58 | 273.78 | 270.85 | 125.32 | 72.83 | 42.98 | 103.21 | 90.47 | 80.71 | 27.81 | 21.38 | 24.38 | 24.97 | 26.83 | 23.02 |
| RTN1 | EFEEDLTGIDDR | 18.58 | 24.03 | 62.20 | 25.43 | 45.87 | 63.74 | NA | 54.73 | 70.19 | 323.46 | 118.15 | 253.66 | 89.08 | 42.36 | 78.64 | 68.90 | 926.66 | 65.03 |
| RXRB | AIILFNPDAK | 219.59 | 260.59 | 292.65 | 217.73 | 245.60 | 247.12 | 94.33 | 59.59 | 32.83 | 84.64 | 67.58 | 77.88 | 17.61 | 15.40 | 17.84 | 18.58 | 6.49 | 19.10 |
| CYP26B1 | VLAVELASTSR | 93.59 | 81.92 | 102.97 | 75.20 | 139.19 | 117.06 | 28.39 | 10.52 | 9.36 | 19.48 | 12.92 | 26.83 | 18.87 | 15.95 | 4.45 | 6.94 | NA | 4.00 |
| RDH10 | ENVYLTAER | 37.37 | 53.42 | 104.06 | 95.95 | 21.93 | 27.75 | 38.54 | 16.33 | 8.62 | 21.68 | 19.83 | 18.13 | 27.54 | 22.90 | 14.75 | 25.02 | 14.61 | 16.59 |
| ALDH1A3 | EEIFGPVQPILK | NA | 175.80 | NA | 68.95 | 1.10 | NA | 31.43 | 23.74 | 15.17 | 29.97 | 30.83 | 27.17 | 12.13 | 14.38 | 19.22 | 19.24 | 14.56 | 17.09 |
| RETSAT | LFPQLEGK | 11.46 | 12.29 | 15.24 | 10.89 | 11.20 | 11.88 | 52.00 | 32.67 | 29.37 | 49.97 | 51.42 | 41.89 | 23.96 | 23.04 | 28.17 | 35.62 | 25.25 | 28.86 |
| RXRG | AIVLFNPDAK | 9.02 | 6.82 | 13.84 | 11.11 | 8.43 | 11.26 | NA | 3.30 | 2.26 | 3.76 | 2.34 | 3.29 | 35.56 | 30.75 | 39.03 | 49.79 | 45.64 | 43.75 |
| CYP26A1 | FSFIPFGGGLR | 19.49 | 23.75 | 26.81 | 20.27 | 18.25 | 12.97 | 4.38 | 2.89 | NA | 2.20 | 2.36 | 1.93 | 2.12 | 1.11 | 1.38 | 1.78 | 3.13 | 1.74 |
| RDH5 | FGLEAFSDSLR | 8.25 | 12.28 | 8.11 | 10.26 | 7.45 | 8.99 | 4.40 | 4.25 | 3.35 | 8.78 | 6.00 | 8.34 | 6.15 | 4.47 | 7.64 | 7.17 | 11.79 | 6.54 |

**Supporting Table S2 : Table of raw protein abundances (fmol/mg protein) across mouse hearts during embryonic, postnatal and adulthood.** Light-to-heavy ratios were calculated using the three most intense fragment ion transitions for each peptide and determined as Σlight/Σheavy. Endogenous peptide abundances were calculated by scaling these ratios to the known heavy peptide input (50 fmol per 5 µg total protein) and are reported as fmol per mg of total protein. Data are shown for embryonic day 9.5 (E9.5), postnatal day 1 (P1), and adult 12-week-old mouse hearts, with n = 6 biological replicates per developmental stage. Values represent unnormalized peptide abundances derived from SureQuant targeted proteomics analysis.

| Peptide Quantification (fmol/mg of protein) | | | | | | | |
|---|---|---|---|---|---|---|---|
| Values scaled for visualization | | | | | | | |
| protein | peptide | rep_1_1 | rep_1_2 | rep_1_3 | rep_14_1 | rep_14_2 | rep_14_3 |
| TTR | TAESGELHGLTTDEK | 29,605.98 | 24,524.29 | 31,237.79 | 26,900.43 | 22,708.51 | 23,021.97 |
| DHRS4 | TALLGLTK | 1,423.28 | 1,193.15 | 1,130.52 | 2,224.82 | 1,530.52 | 1,608.47 |
| RBP4 | FSGLWYAIAK | 985.37 | 1,227.04 | 1,055.52 | 939.37 | 503.95 | 482.75 |
| RDH14 | TVLITGANSGLGR | 910.30 | 918.71 | 829.14 | 1,029.98 | 674.58 | 623.79 |
| ALDH1A1 | QAFQIGSPWR | 358.04 | 334.85 | 212.62 | 801.74 | 536.39 | 654.44 |
| RDH14 | ELDLASLR | 354.59 | 350.15 | 308.76 | 504.94 | 347.37 | 362.37 |
| RETSAT | LFPQLEGK | 355.41 | 443.12 | 75.69 | 488.94 | 241.28 | 229.10 |
| RDH13 | TVIVTGANTGIGK | 257.74 | 254.48 | 222.73 | 427.41 | 288.28 | 325.09 |
| ALDH1A2 | EEIFGPVQEILR | 264.45 | 239.83 | 220.28 | 144.15 | 41.79 | 83.72 |
| ALDH1A3 | EEIFGPVQPILK | 102.69 | 81.89 | 52.03 | 108.03 | 74.92 | 77.86 |
| RXRB | AIILFNPDAK | 68.18 | 65.56 | 53.13 | 71.08 | 43.06 | 42.71 |
| RXRA | AIVLFNPDSK | 52.52 | 49.58 | 46.31 | 65.87 | 39.59 | 38.44 |
| RXRG | AIVLFNPDAK | 27.93 | 22.12 | 23.53 | 63.07 | 46.06 | 45.49 |
| RDH8 | DLYLPASR | 17.67 | NA | NA | 17.48 | 17.28 | 16.00 |
| RDH10 | LFALEFAR | 8.90 | 10.23 | 8.99 | 13.71 | 8.62 | 9.15 |
| RDH12 | LANVLFTR | 3.23 | 2.30 | 2.04 | 5.34 | 4.31 | 4.32 |
| RDH5 | FGLEAFSDSLR | 3.40 | 3.76 | 2.44 | NA | NA | NA |

**Supporting Table S3 : Table of raw protein abundances (fmol/mg protein) across postnatal days 1 and 14**. Light-to-heavy ratios were calculated using the three most intense fragment ion transitions for each peptide and determined as Σlight/Σheavy. Endogenous peptide abundances were calculated by scaling these ratios to the known heavy peptide input (50 fmol per 5 µg total protein) and are reported as fmol per mg of total protein. Data are shown for each developmental stage with n = 3 biological replicates per time point. Values represent unnormalized peptide abundances obtained by SureQuant targeted proteomics analysis.

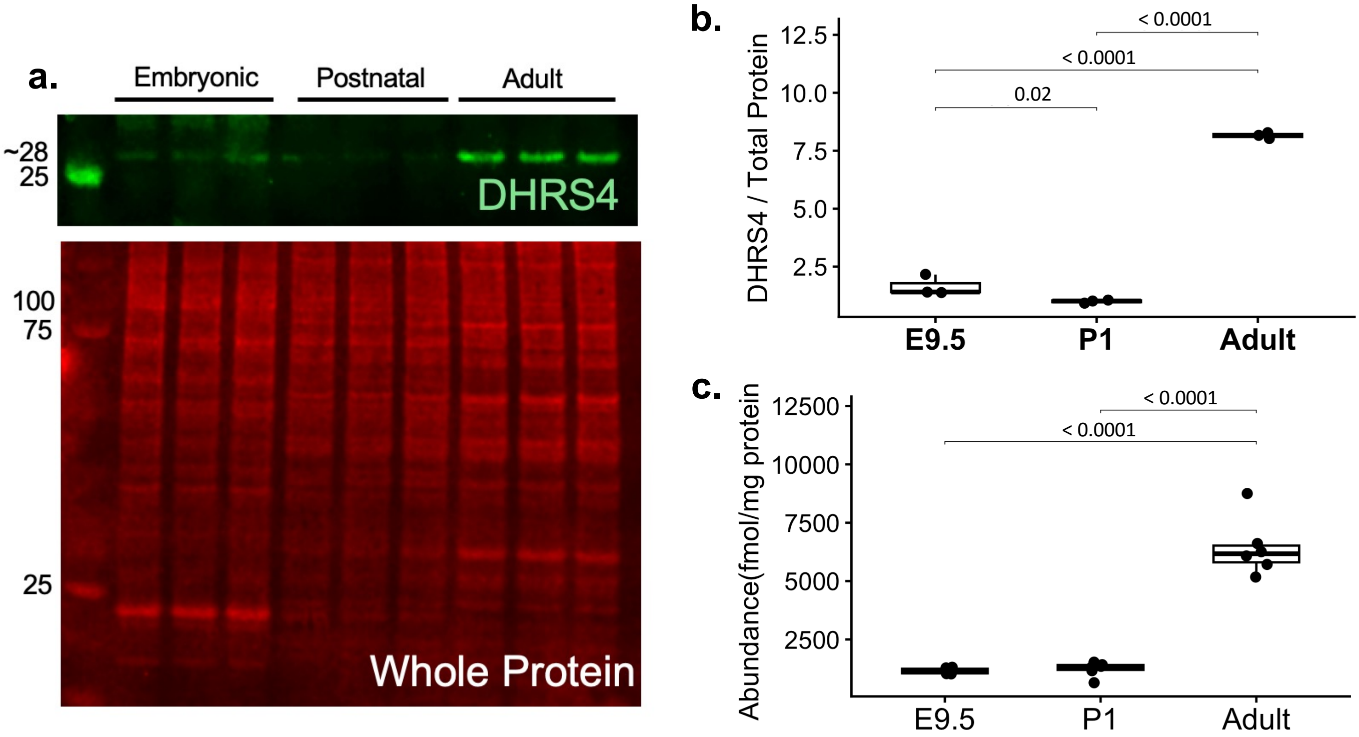


**Supporting Figure S3. Validation of developmental changes in retinoid pathway proteins in mouse hearts at embryonic day 9.5 (E9.5), postnatal day 1 (P1), and adult (12 weeks) by Western blotting.** Western blot analyses of DHRS4 showed trends consistent with those observed by SureQuant PRM analysis. Mouse heart samples were analyzed with $n$ = 3 biological replicates per developmental stage for Western blotting and $n$ = 6 biological replicates per developmental stage for SureQuant PRM analysis. (a) Representative Western blot for DHRS4 showing an immunoreactive band at approximately 28 kDa (b) shows protein abundance determined by Western blotting and normalized to total protein staining. (c) shows protein abundance measured by SureQuant PRM and reported as fmol/mg of total protein, and panel Data are presented as mean ± SD. Statistical significance between developmental stages was assessed using one-way ANOVA followed by pairwise post hoc comparisons. Significant pairwise differences are indicated on the plots ($p < 0.05$).

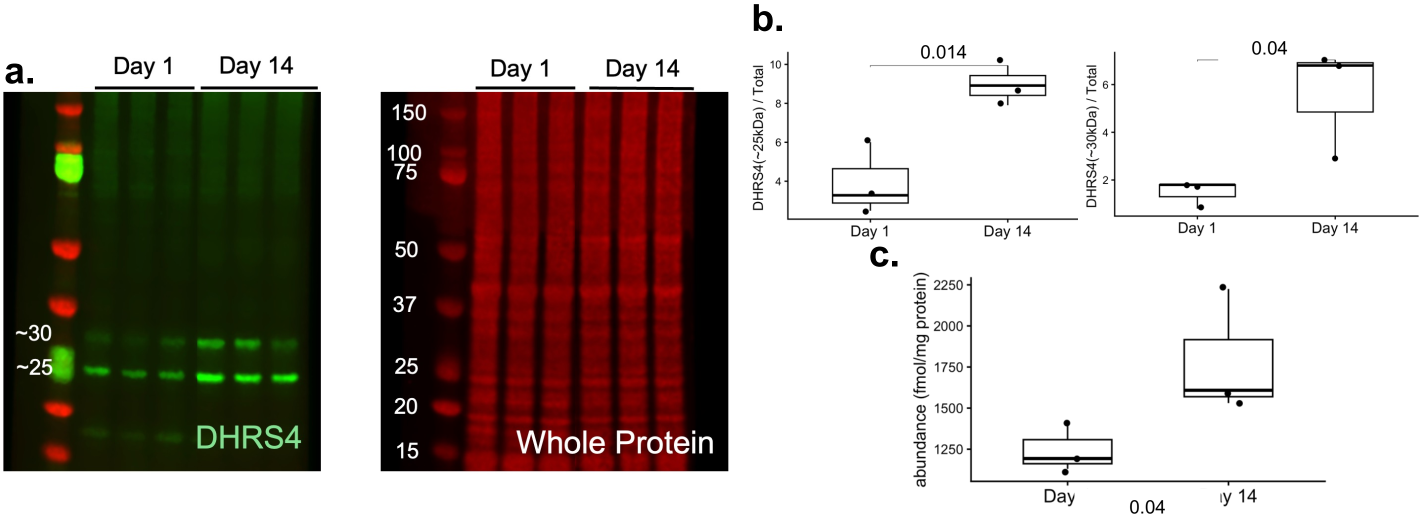


**Supporting Figure S4. Validation of developmental changes in retinoid pathway proteins in rat hearts at postnatal day 1 (P1) and postnatal day 14 (P14) by Western blotting.** Western blot analyses of DHRS4 showed trends consistent with those observed by SureQuant PRM analysis. Rat heart samples were analyzed at P1 and P14 (*n* = 3 biological replicates per time point). (a) Representative Western blot showing two immunoreactive bands at approximately 25 and 30 kDa, which may represent alternative isoforms, post-translationally processed forms, or proteolytic cleavage products. (b) shows protein abundance determined by Western blotting and normalized to total protein staining. (c) shows protein abundance measured by SureQuant PRM and reported as fmol/mg of total protein, and panel Data are presented as mean ± SD. Statistical significance between P1 and P14 groups was assessed using an unpaired two-sided t-test. Significant differences are indicated on the plots ($p < 0.05$).
